# Short-term alterations in dietary amino acids override host genetic susceptibility and reveal mechanisms of *Salmonella* Typhimurium small intestine colonization

**DOI:** 10.1101/2025.03.25.645332

**Authors:** Nicolas G. Shealy, Madi Baltagulov, Camila de Brito, Anna McGovern, Pollyana Castro, Alexandra C. Schrimpe-Rutledge, Clara Malekshahi, Simona G. Condreanu, Stacy D. Sherrod, Somnath Jana, Katerina Jones, Tamara Machado Ribeiro, John A. McLean, Daniel P. Beiting, Mariana X. Byndloss

## Abstract

In addition to individual genetics, environmental factors (e.g., dietary changes) may influence host susceptibility to gastrointestinal infection through unknown mechanisms. Herein, we developed a model in which CBA/J mice, a genetically resistant strain that tolerates intestinal colonization by the enteric pathogen *Salmonella* Typhimurium (*S.* Tm), rapidly succumb to infection after exposure to a diet rich in L-amino acids (AA). In mice, *S.* Tm-gastroenteritis is restricted to the large intestine (cecum), limiting their use to understand *S*. Tm small intestine (ileum) colonization, a feature of human Salmonellosis. Surprisingly, CBA mice fed AA diet developed ileitis with enhanced *S*. Tm ileal colonization. Using germ-free mice and ileal-fecal slurry transplant, we found diet-mediated *S*. Tm ileal expansion to be microbiota-dependent. Mechanistically, *S*. Tm relied on Fructosyl-asparagine utilization to expand in the ileum during infection. We demonstrate how AA diet overrides host genetics by altering the gut microbiota’s ability to prevent *S.* Tm ileal colonization.

## INTRODUCTION

The WHO estimates that 33 million individuals (1 in 10) worldwide are impacted by foodborne gastroenteritis each year^1^. One of the major causes of such illnesses is the bacterium *Salmonella enterica* serovar Typhimurium (*S.* Tm)^2^. Interestingly, 80% of all *S.* Tm infections have no known explanation for host susceptibility^2^. While it is known that a history of antibiotic and steroid use can influence *S.* Tm-induced gastroenteritis, it remains to be fully elucidated how environmental factors such as diet influence host susceptibility and pathogen virulence.

Multiple genome-wide association studies (GWAS) have identified genetic factors that impact the host’s ability to control bacterial infections, highlighting the critical role of genetic-based predisposition in infectious disease severity and outcomes^3^. For instance, mutations in the metal-ion transporter in professional phagocytes encoded by *NRAMP1* have been implicated in the resistance to intracellular bacterial pathogens (e.g., *Mycobacterium tuberculosis* and *Salmonella enterica*)^3^. Indeed, mice expressing a non-functional *Nramp1* allele, such as C57Bl6, present with severe and fatal typhoid-like infections^4^. It remains unknown, however, how the environment influences genetic-based resistance mechanisms to bacterial infections. Diet, smoking, pollution, and other environmental co-morbidities have been shown to exacerbate host susceptibility despite host genetics^5^, through largely unknown mechanisms. Such findings implicate the role of environmental factors in modulating genetic susceptibility to infectious disease. An underappreciated aspect linking environmental factors and genetic-based susceptibility to bacterial infections is the impact of dietary habits on the gut microbiota, the resident collection of commensal bacteria that provide colonization resistance against enteric pathogens^6–14^. Specifically, many investigations of the role of diet have focused on the contribution of dietary carbohydrates and fat and disregarded protein despite it being a key macronutrient. The contribution of dietary protein sources and their influence on susceptibility to enteric pathogens such as *S.* Tm remains to be elucidated.

The small intestine represents a pertinent gut geography for host-pathogen interactions^15^. Comprising much of the gastrointestinal tract by length, the small intestine is the primary location of nutrient absorption and the focal site of *S*. Tm-induced gastroenteritis in humans^16^. Despite *S.* Tm causing small intestinal disease in humans, few tractable animal models exist to address the dynamics of *S.* Tm colonization of the small intestinal lumen. While animal models exist for studying *S*. Tm small intestine pathogenesis, such as ligated ileal loop models in calves, guinea pigs, and chickens^17^, they show significant limitations, ranging from high-cost genetic manipulation to low physiological relevance to humans^17,18^. Murine models of infection, the most genetically tractable option, usually develop *S.* Tm-induced intestinal inflammation in the proximal portion of the large intestine, the cecum, a region morphologically distinct from the human anatomical homolog ^16^. It has been shown that the large and small intestines significantly differ in biotic and abiotic factors, such as the community of gut microbes, pH, and nutrient sources in each gut geography^19^. Thus, there is a pressing need to develop easily accessible animal models to study mechanisms used by *S.* Tm to gain access to nutrients in the small intestine during infection.

Herein, we show that alterations in dietary amino acid influence the susceptibility of genetically resistant CBA/J mice to infection by *S.* Tm. Interestingly, increased susceptibility is not due to the overall inability of the murine host to control infection (e.g., exacerbated susceptibility to systemic infection) or differences in host small intestine transcriptome prior to infection. Instead, significant reductions in microbial diversity of the small intestine impact the ability of the small intestine microbiota to prevent *S.* Tm colonization and expansion. Lastly, using this model, we identified small intestine-relevant metabolic pathways necessary for *S*. Tm expansion, such as using Fructosyl-Asparagine, which enables expansion in the ileum.

## RESULTS

### Mice fed a free L-amino acid diet demonstrate marked susceptibility to oral challenge with *S*. Tm

Environmental factors such as diet play a key role in the composition and function of the gut microbiota and the susceptibility to enteric pathogens such as *Salmonella enterica* serovar Typhimurium (*S.* Tm)^20^. While diet has been shown to influence the gut microbiota and *S.* Tm infection, published studies focus primarily on the impact of dietary carbohydrates (e.g., fiber or high sugar) or fats^20^, and ignore other significant diet constituents such as protein and amino acids. Indeed, while intermediate protein digestion products such as di- and tripeptides are known to influence host digestion, the local gut immune compartment, as well as the gut microbiota, little is known about their role during infection with enteric bacterial pathogens^21–23^. To address this knowledge gap, we sourced a diet that supplied free L-amino acids (AA diet) and was calorically matched to the chow used in our vivarium. We chose to use free amino acids rather than alternative protein sources to control the distribution of amino acids supplied to the mice between diets (**Table S1**). Notably, the only difference between diets was the form in which the amino acids were given, either via whole protein casein or as L-amino acids in a matching distribution to that found in casein. To interrogate the role of dietary amino acids during *S.* Tm infection, we chose to leverage the genetically resistant mouse background, CBA/J, that does not require streptomycin pre-treatment for induction of *S*. Tm-mediated intestinal inflammation, which could obscure diet-dependent alterations. CBA/J mice can restrict *S*. Tm, developing gastroenteritis between seven to ten days post-infection (d.p.i.), with mice usually surviving the infection^24^.

To test the impact of dietary amino acids on *S.* Tm-induced gastroenteritis, CBA/J mice were switched to an AA diet two days before oral challenge with 10^7^ colony-forming units (CFUs) *S*. Tm (or were maintained on chow) and mice were allowed to carry the pathogen for up to seven d.p.i. (**Fig. 1A**). We chose an infectious dose of 10^7^ CFU, as to our surprise AA diet-fed mice infected with the conventional dose of 10^9^ CFU demonstrated severe disease within three to five d.p.i. (data not shown). Additionally, we chose the two-day time point for diet exposure before infection to model short-term alterations in diet and prevent marked diet alterations to host physiology and immune responses before infection. *S.* Tm-infected mice fed an AA diet experienced more severe disease than chow-fed mice, characterized by significant weight loss by five d.p.i. (**Fig. 1B**), and significantly higher *S*. Tm burden in feces as early as one d.p.i. (**Fig. S1A)** that lasted throughout the infection (**Fig. 1C**). Taken together, these data suggest that an AA diet accelerates gastrointestinal disease onset following the oral challenge of CBA/J mice with *S*. Tm with increased burden and mortality.

**Figure 1:**
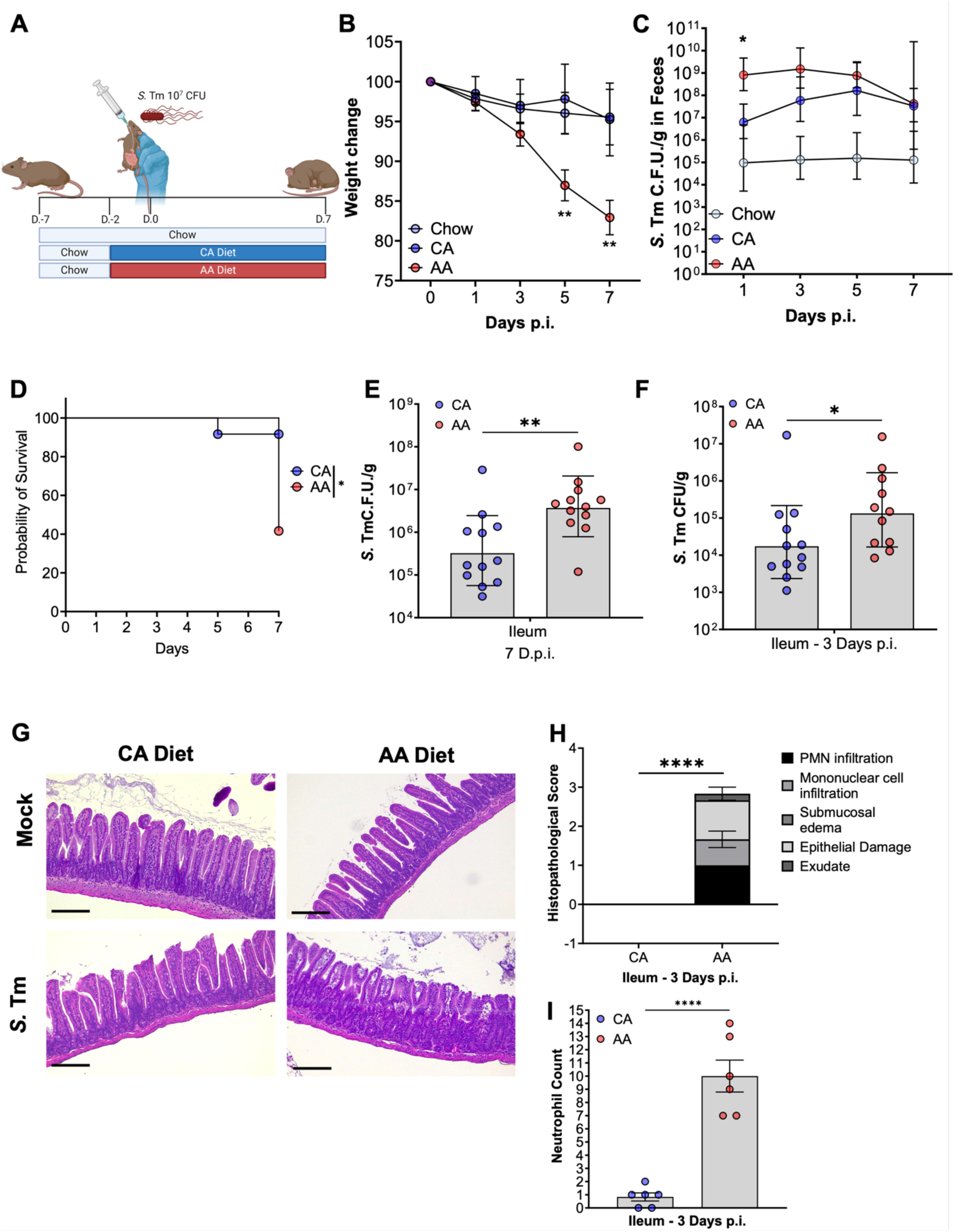
AA diet-fed CBA/J mice demonstrate increased fecal and ileal *S*. Tm burden associated with mild to moderate ileitis and mortality. (A-E) (A) Infection is schematic: female CBA/J mice were either maintained on vivarium chow or switched to a company-matched whole protein casein diet (CA) or L-amino acid diet (AA) two days before oral challenge with *S*. Tm 10^7^ C.F.U. and maintained for up to 7 days p.i. (B) Weight lost post-infection (C) Fecal burden (D) Survival (E) Ileal burden 7 days p.i (F) Ileal burden 3 days p.i. (G) Representative images of H&E ileal tissue from 0 and 3 days p.i. (H) Histopathology scoring of ileal tissue. Bar = 200 µm. (I) Neutrophil count from (G-H). Geometric mean and geometric SD, (B-C) Two-way ANOVA, (D) Log-rank (Mantel-Cox) Test, E-F) student’s t-test, (H-I) Mann-Whitney; N=6-10 per diet; P=*, <0.05; **, <0.01; ****, <0.0001.

Other groups have shown that variations in manufacturing practices of rodent diets between companies, such as the incorporation of heavy metals and phytoestrogens, can alter host physiology^25^. To address the potential of company practices confounding differences in *S*. Tm susceptibility of CBA/J mice, we sourced an additional diet from the same company as the AA diet and replaced the free L-amino acids with casein (CA diet) to recapitulate nutrients found in our vivarium chow. All other components of the diets were maintained, including the addition of sodium bicarbonate to compensate for the high amino acid concentration of our AA diet (**Table S1).** As previously performed, mice were switched to either an AA or a CA diet two days before the *S.* Tm challenge. In line with previous results using chow as a control, mice switched to an AA diet lost considerably more weight than those fed a CA diet starting at five d.p.i. (**Fig. 1B**), had increase *S*. Tm fecal burden during early infection (**Fig. 1C**) and harbored significantly more *S*. Tm in ileum content at three and seven d.p.i. (**Fig. 1E-F**). In line with the increased *S*. Tm burden, AA diet-fed mice demonstrate accelerated disease onset and mortality, resulting in a loss of ∼60% of the population as compared to only a ∼10% loss in CA-die fed mice (**Fig.1D)**. Additionally, AA diet-fed mice demonstrated significantly increased inflammation characterized by mild epithelial damage and neutrophil influx in the ileum compared to CA diet-fed mice three d.p.i. (**Fig. 1G-I**). Similar results were found in the cecum as well as the proximal colon of AA diet (**Fig. S1F-G**). Importantly, AA diet alone did not cause alterations in uninfected mice weight (**Fig. S1B**), food consumption (**Fig. S1C**), gut morphology (**Fig. 1G**), or luminal pH (**Fig. S1D**). Our data demonstrates that diet containing casein as the protein source (chow and CA diet) protect genetically resistant mice from lethal *S.* Tm gastroenteritis, while L-amino acid-based diet (AA diet) promotes susceptibility to *S.* Tm infection through unknown mechanisms. Importantly, we decided to use diets from the same vendor (CA and AA diets) for our additional studies to avoid the impact of protein source-independent factors related to manufacturing practices on our phenotypes.

Interestingly, the *S.* Tm fecal burden (proxy for large intestine colonization) in CA and AA diet-fed mice was comparable by seven d.p.i. (**Fig. 1C**) despite major differences in disease manifestation and weight loss (**Fig. 1B-D**). Additionally, no differences were observed in *S.* Tm colonic burden between CA and AA diet-fed mice, and both groups harbored significantly more *S*. Tm in their colon content than chow-fed mice (data not shown). While *S*. Tm abundance in the colon of CA and AA diet-fed mice at seven d.p.i. was equivalent, the same was not true for other segments of the intestinal tract. AA diet-fed mice demonstrated a significantly increased *S.* Tm burden in ileal content at seven d.p.i., suggesting that AA-diet feeding enabled *S*. Tm marked colonization of the distal small intestine (**Fig. 1E**). Thus, we interrogated how early during infection *S*. Tm could colonize and expand in the ileum. To do so, we performed the experiment as previously described (**Fig. 1A**) and assessed *S.* Tm intestinal colonization at three d.p.i., before significant differences in weight loss between experimental groups occur (**Fig. 1B**). To our surprise, as early as three d.p.i., AA diet-fed mice harbored significantly more *S*. Tm in the lumen of the ileum than CA diet-fed mice (**Fig. 1F**) which was associated with a mild to moderate ileitis (**Fig. 1G-H**) and mild neutrophil recruitment to the ileal mucosa (**Fig. 1I**). The literature demonstrates that CBA/J mice typically develop *S.* Tm-enterocolitis focally located in the cecum^26,27^. However, mice fed an AA diet developed disease starting in the ileum (**Fig. 1E-I**), an intestinal geography highly relevant to human manifestations of Salmonellosis^17,28,29^. Thus, our results suggest that the AA diet enables *S*. Tm colonization of the distant small intestine, allowing for the study of *S*. Tm pathogenesis in the intestinal sites more physiologically comparable in geography to *S.* Tm-induced gastroenteritis in humans. Using our AA diet model, we continued to explore the early three-day time point to understand the important factors for the colonization and expansion of *S*. Tm in the ileum.

### Mice fed a free L-amino acid diet do not have increased susceptibility to systemic *S*. Tm infection

Although *S.* Tm is commonly transmitted through the fecal-oral route, this pathogen is also able to cause systemic infections and replicates within the peritoneal cavity, liver, and spleen^17^. Transmission to the liver and spleen is significantly impacted by the bottleneck that exists through oral gavage infections due to the acidic pH of the stomach and the acid sensitivity of *S*. Tm^26,27^. To overcome the gut-intrinsic bottleneck and directly interrogate the ability of *S.* Tm to survive within systemic sites, we infect mice directly via intraperitoneal injection (IP) as is commonly done for models of systemic *Salmonella* serovars such as Typhi and Paratyphi^30^. Importantly, the severity of disease following IP is directly impacted by the ability of the host immune system to restrict pathogen replication in systemic organs and bypass the bottlenecks associated with intragastric infection^17,28^.

One possible explanation for the increased susceptibility of AA diet-fed CBA/J to *S.* Tm infection would be that the AA diet was disrupting the ability of the host to control *S.* Tm systemic proliferation. Thus, to interrogate if an AA diet altered the ability of CBA/J to handle systemic challenges with *S*. Tm, we altered the route of infection from per oral to IP route with 10^4^ CFU and assessed *S.* Tm burden at four d.p.i. (**Fig. 2A)**. No significant differences were found in the *S*. Tm burden of the intraperitoneal cavity (**Fig 2B**) nor systemic dissemination sites such as the liver (**Fig. 2C**) and spleen (**Fig. 2D**) between different diets. One major protective factor influencing CBA/J susceptibility to *S*. Tm infection is their expression of the functional *Nramp1* allele by professional phagocytes such as macrophages^3,4^. Natural-resistance-associated macrophage protein 1 or NRAMP1 enables professional phagocytes to effectively kill *S.* Tm by leveraging oxidative stress-mediated killing through the sequestration of divalent ions, limiting systemic spread ^3,4^. To further interrogate the necessary component of CBA/J to control the systemic spread of *S*. Tm, we performed qPCR on ileal tissue from orally infected mice fed either a CA or AA diet at three d.p.i.. We found no significant defects in the expression of *Nramp1* (**Fig. S2**). Given our findings through IP infection as well as qPCR of *NRAMP1* in the ileum of *S*. Tm infected mice, we reasoned there were no alterations in the ability of AA diet-fed mice to control systemic *S*. Tm as compared to either Chow-maintained or CA diet-fed mice, and we hypothesized the mechanism of susceptibility to be independent of *NRAMP1*, and restricted to diet-mediated changes to the host’s gastrointestinal tract.

**Figure 2:**
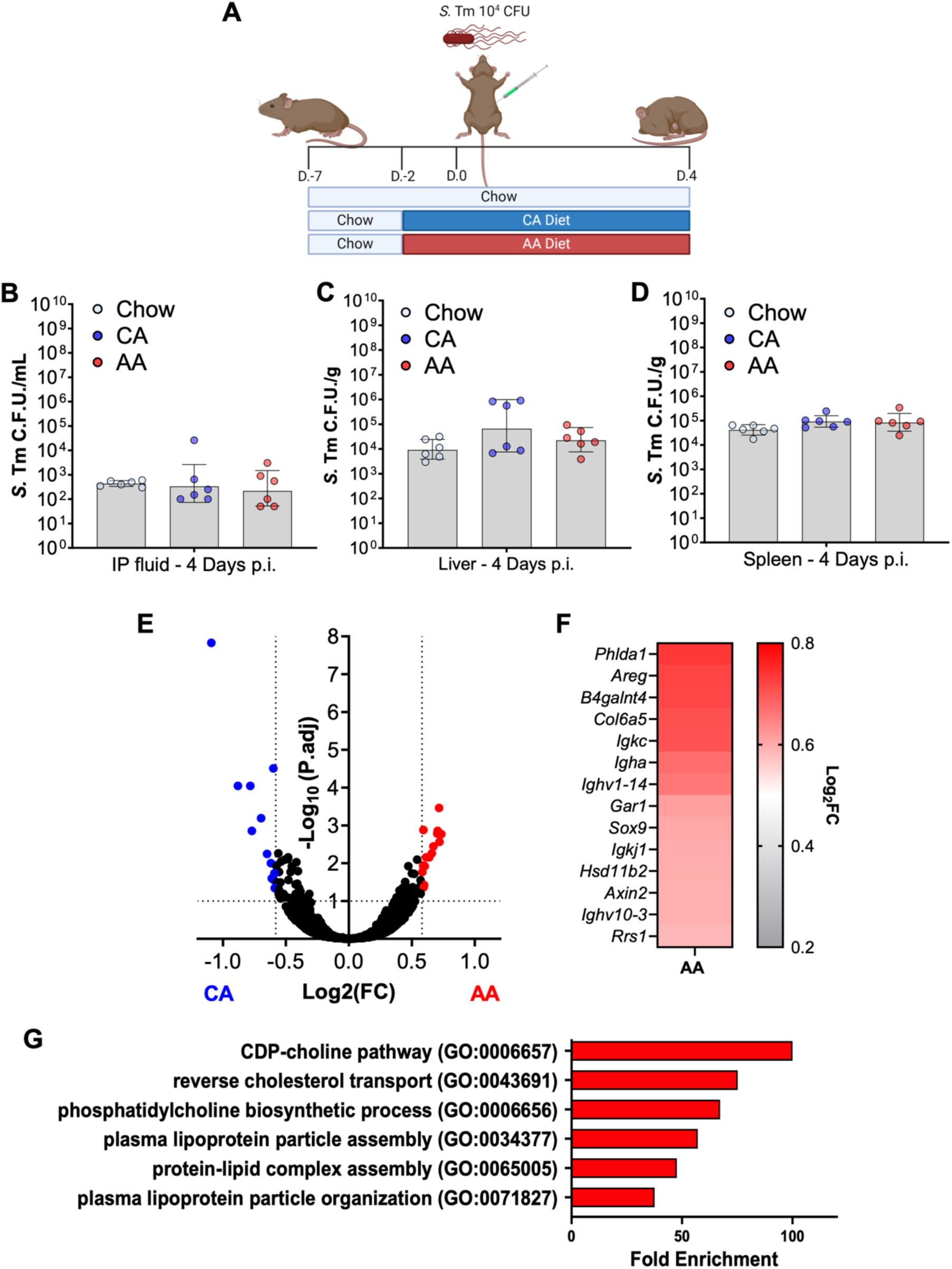
AA diet does not alter host ability to handle *Salmonella* Typhimurium systemic infection, nor does it significantly alter ileum transcriptomic profile. (A-D) (A) Infection is schematic: female CBA/J mice were either maintained on vivarium chow or switched to a company-matched whole protein casein diet (CA) or L-amino acid diet (AA) two days before intraperitoneal (IP) challenge with *S*. Tm 10^4^ C.F.U. and maintained up to four days p.i. (B-D) *S*. Tm burden in (B) liver, (C) spleen, and (D) IP lavage fluid. (E-F) RNAseq from bulk ileal tissue of uninfected AA or CA diet-fed mice. N=3 per condition. Volcano plot (E) and Heatmap (F) of upregulated genes in the of ileum of AA diet fed uninfected mice. (G) Gene ontology of unique genes upregulated in the ileum of uninfected mice following the switch to AA as compared to CA using the Panther algorithm. (B-D) Geometric mean and geometric SD, N=6-10 per diet.

### AA diet induces mild changes to the ileum mucosa before infection

Given that an AA diet did not alter the host’s susceptibility to systemic *S.* Tm infection, we postulated that an AA diet disrupted the ileal mucosa protective functions to enable increased *S.* Tm intestinal colonization, as alterations in barrier function, such as the loss of beta-catenin or zonula occludens, have been shown to contribute to *S.* Tm gastroenteritis^31^. To further interrogate the impact of AA diet on the gastrointestinal tract, we performed bulk RNA sequencing (RNA-seq) of the ileal tissue from mice exposed to either CA or AA diet for two days, before infection (**Fig. 2E-F**). To our surprise, we found only a limited number of significantly differentially expressed genes (DEGs), 25 DEGs between diets in the ileum, with most genes differentially regulated being implicated in the development and proliferation of the epithelium of the small intestine, such as associations with Wnt signaling. Wnt signaling is essential for maintaining barrier function in the gut epithelium and is associated with gastrointestinal diseases such as colitis and colorectal cancer^32^. Specifically, we observed an increased expression in proliferation markers such as *Axin2* and *Areg*^33–35^. Counterintuitively, we also observed a significant increase in *Sox9* and *Lbh,* which regulate Wnt signaling in the gut^36^. Unrelated to Wnt, we observed a significant increase in the glycosyltransferase, *B4galnt4*, which has been implicated in alterations in the glycosylation of extracellular matrix proteins and, more importantly, mucus that forms the protective barrier between the gut epithelium and the microbiota and promotes and tolerogenic relationship between the two^37^. Lastly, we note a significant increase in the expression of B cell-associated genes such as the production of immunoglobulins via *Igha*, *Ighv1-14*, *Igkj1*, and *Igkc*, representing one of the main functions of B cells in the gut^38^.

While we observed only a few DEGs between CA and AA diets before infection, we also wanted to investigate whether changing the mouse diet from chow to our experimental diets impacted small intestine physiology. To do so, we compared the transcriptomic profile of mice maintained on Chow to those switched to a CA or AA diet and then subsequently compared the enriched genes between diets. From these DEGs, we then interrogated those enriched in AA diet by performing a gene ontology enrichment analysis (GO). We found a significant association with genes involved in fat and cholesterol metabolism in AA diet-fed mice (**Fig. 2G**). Interestingly, the small intestine represents the site of fat and cholesterol absorption. Altogether, these data suggest that an AA diet impacts proliferation and potential cell populations within the gut epithelium and associated lymphoid tissues, such as Peyer’s patches, that *S*. Tm commonly targets during the infection cascade^17,39^. Our transcriptomics data also demonstrates that the effect of AA diet in the ileum physiology before infection is limited, with few significant DEGs found between diets, and with none of them having over a two-fold induction in the ileum of AA diet-fed mice. Thus, our results suggest that changes to ileal physiology before infection may not be the main reason for the increased susceptibility of AA diet-fed mice to *S*. Tm ileal colonization. Thus, we chose to explore other factors that could explain the phenotype such as the diet’s impact on the gut microbiota.

### The susceptibility of AA diet-fed mice to *S.* Tm infection is microbiota-dependent

Published studies on the effect of diet on the microbiota demonstrate that microbiota alterations occur as quickly as 24 hours post-diet alteration and that diet-induced alterations directly influence susceptibility to enteric pathogens by altering colonization resistance and the host response^40^. The germ-free mouse model enables the direct interrogation of host factors that contribute to the host’s susceptibility to enteric pathogens in the absence of a resident gut microbiota^41^. To further interrogate the AA diet-mediated alterations in host susceptibility to colonization of *S.* Tm and severe disease, independent of the gut microbiota, we repeated our AA diet-fed murine *S.* Tm infection model in a Germ-Free Swiss Webster (GF-SW) mouse background to determine what gut intrinsic alterations occurred due to switching mice to an AA diet. GF-SW mice like CBA/J are genetically resistant to *S*. Tm and develop gastroenteritis without antibiotic pre-treatment^41^. GF-SW mice orally infected with *S*. Tm harbored a high pathogen burden in the ileum independent of diet background (**Fig. 3A**). Together, these findings suggest that AA diet does not promote *S*. Tm’s small intestine colonization in the absence of a gut microbiota.

**Figure 3:**
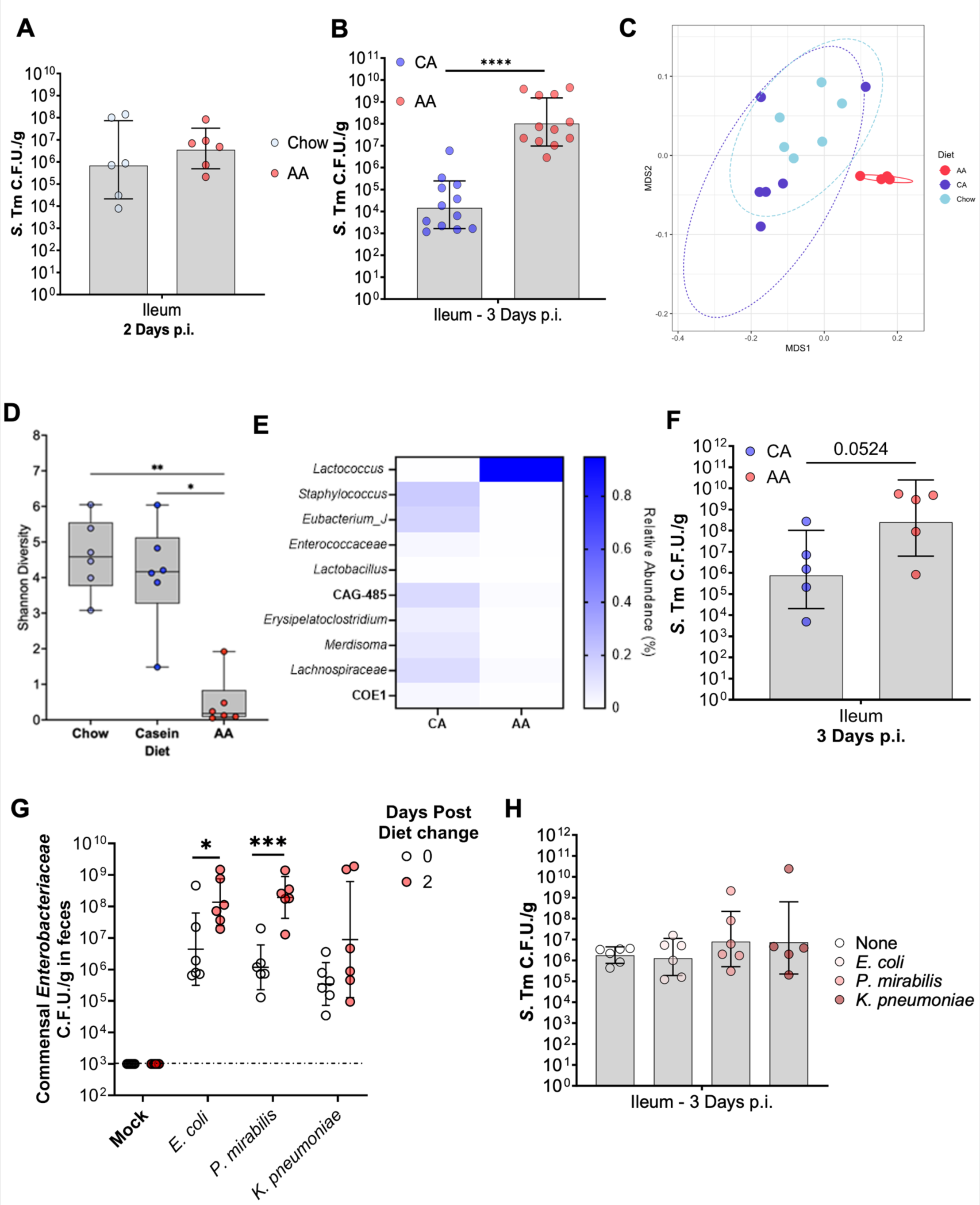
AA diet significantly alters the composition of the gut microbiota and disables the ability of commensal *Enterobacteriaceae* to confer resistance to *S*. Tm. (A) Germ-free Swiss Webster maintained on vivarium Chow or switched to AA-diet two days prior to infection with *S.* Tm and carried for two days post-infection. (B) Terminal ileal burden of *S.* Tm from GF-SW mice received ileal-fecal transplant five days before the diet switch from either CA or AA diet-fed. Two days before infection, mice switched to either a CA or AA diet and were subsequently infected with *S.* Tm and carried for 3 days before euthanasia. N=10-12 per group. (C) Nonmetric multidimensional scaling plot comparing beta diversity via weighted unifrac between diet groups. (D) Alpha diversity of 16rDNA sequencing from the ileal contents of uninfected mice maintained on chow or switched to either a CA or AA diet for two days before euthanasia. (E) Heatmap of top ten most abundant genera in the terminal ileum, the relative abundance of each genus in CA or AA diet-fed mice two days following diet switch without infection. (F) Terminal ileal burden of *S.* Tm from GF-SW pre-colonized with a 13-member defined microbiota (see Materials & Methods) 5 days before diet switch to either a CA or AA diet and subsequent infection with *S.* Tm. (G-H) Conventional CBA mice from Jackson Labs were colonized with representative commensal *Enterobacteriaceae* or mock control, five days before being switched to an AA diet and subsequent infection with *S.* Tm. (G) Commensal *Enterobacteriaceae* burden in feces before and after diet switch. (H) *S.* Tm burden in the terminal ileum three days post-infection following euthanasia. N=6, Multiple t-test or One-way ANOVA, N = 4-6 per group P=*, <0.05; **, <0.01; ***, <0.001; ****, <0.0001.

To interrogate the gut microbiota’s role in the susceptibility of AA diet-fed mice, we performed an ileal-fecal microbiota transplant (IFMT) from uninfected CBA/J fed either a CA or AA diet for two days by combining the content of both small and large intestines. CA- or AA-IFMTs were then implanted into GF-SW mice via oral gavage. After allowing the resultant conventionalized SW mice and the microbiota to stabilize for five days following transplantation, we challenged IFMT-recipient mice with *S*. Tm via intragastric infection, as we had previously performed (**Fig. 1A**). While mice given an IFMT from casein-fed donors were resistant to ileal colonization of *S*. Tm, littermates who received an IFMT from AA diet-fed mice demonstrated significantly higher ileal burden at three days post-infection (**Fig. 3B**). Altogether, these data suggest the susceptibility of AA diet-fed mice to *S.* Tm gastrointestinal infection is microbiota-dependent.

### AA diet-mediated changes in gut microbiota composition promote *S*. Tm small intestine colonization

Our findings suggested that AA-diet-mediated changes to the small intestine microbiota contribute to *S*. Tm’s ability to colonize the small intestine of AA-fed mice. To interrogate how an AA diet influenced the gut microbiota structure of the distal small intestine, we performed 16S rRNA sequencing of ileal content from mice switched to either a CA or AA diet for two days before sample collection prior to infection. We first observed that the microbial profile of AA diet clustered away from that of both Chow and CA fed mice, which clustered together via weighted Unifrac (**Fig. 3C**). Additionally, compared to both Chow-maintained and mice switched to a CA diet, the microbiota of AA diet-fed mice demonstrated a significant reduction in alpha diversity, a metric of species richness (**Fig. 3D**). When assessing the differences between microbiota composition at the genus taxonomic level, AA diet-fed mice demonstrate a significant decline in the relative abundance in nine of the ten major genera of ileal microbiota, with an overrepresentation of a single genus, *Lactococcus,* in comparison to other diet backgrounds (**Fig. 3E**).

To interrogate whether the AA-mediated increase in *Lactococcus* abundance in the small intestine microbiota was necessary to *S*. Tm’s ability to colonize the small intestine of AA-fed mice, we repeated the GF-SW infection, now colonizing GF-SW mice with representatives of the nine major genera (*Staphylococcus, Eubacterium, Enterococcus, Lactobacillus, CAG-485, Eyrispelatoclostridium, Merdisoma, Lachnospiraceae,* and *COE-1*). Importantly, we withheld any *Lactococcus* representatives from this defined microbiota. After the microbiota stabilization, mice were switched to a CA or AA diet for two days as previously performed and intragastrically infected with *S*. Tm. AA diet-fed gnotobiotic mice colonized with a defined microbiota demonstrated a significantly higher *S*. Tm burden in the ileum compared to CA diet-fed littermates three d.p.i. (**Fig. 3F**). This data suggests that AA diet-dependent loss of microbial diversity rather than the abundance of *Lactococcus* in the distal small intestine microbiota influences colonization of the ileum.

Work by Osbelt and Eberl *et al.* demonstrates that one of the major protective functions of the microbiota is nutrient sequestration, which fortifies the ability of commensal *Enterobacteriaceae* to compete with invading pathogens for the remaining nutrients niches^20,42,43^. All our experiments are performed on mice from Jackson labs, which lack commensal *Enterobacteriaceae*, which are key commensal microbes providing colonization resistance to *S.* Tm in the large intestine^44^. To interrogate if the microbiota of AA diet-fed mice was able to protect the host from *S.* Tm infection in the presence of the commensal *Enterobacteriaceae*, we pre-colonized CBA/J mice from Jackson laboratories with one of three commensal *Enterobacteriaceae* shown to confer colonization resistance against *S*. Tm: *Escherichia coli*, *Klebsiella pneumoniae*, or *Proteus mirabilis*^20,42–44^. After pre-colonization, mice were allowed to acclimate for five days before diet switch to our AA diet as previously performed. Interestingly, all pre-colonized mice fed the AA diet demonstrated an increased fecal burden of commensal *Enterobacteriaceae,* specifically *E. coli* and *P. mirabilis,* but not *K. pneumoniae* upon diet switching before infection (**Fig. 3G**). However, following infection at three d.p.i., we detect no significant differences in the burden of the *S.* Tm in the ileum of AA diet fed mice, independent of commensal *Enterobacteriaceae* colonization status, suggesting that *Enterobacteriaceae* do not prevent *S.* Tm ileal colonization in the AA diet background (**Fig. 3H**). Together, these data suggest that potentially the nutrient sources fueling *S*. Tm expansion in the ileum of AA diet-fed mice are either not used by these commensal *Enterobacteriaceae* representatives, or that the changes in the gut microbiota of AA diet-fed mice alter the metabolic profile of the intestinal lumen in a way that opens possible niches to accommodate both commensal *Enterobacteriaceae* and *S.* Tm without promoting competition before the onset of severe inflammation.

### *S*. Tm does not require respiration nor the activity of its two T3SS to colonize the ileum of AA diet-fed mice

To invade and replicate within intestinal epithelial cells, *S.* Tm requires the activity of its two type-III secretion machinery and the associated effectors encoded on the *Salmonella* Pathogenicity Islands 1 and 2 (SPI-1, SPI-2, respectively) (**Fig. 4A**)^45,46^. Indeed, it is known that the deletion of one or both T3SS results in a decreased ability of *S.* Tm to invade and cause gastrointestinal disease. To determine the role of the two T3SSs in the AA diet susceptibility model, we repeated our infection of AA diet-fed mice with either a virulent wild-type strain or a strain lacking the activity of both T3SSs (Δ*invA*Δ*spiB*). While AA diet fed mice lost significantly more weight than CA diet-fed littermates, 3 d.p.i. when infected with wild-type (WT) *S*. Tm, no significant differences were seen in weight loss between diets when mice were infected with the avirulent Δ*invA*Δ*spiB S.* Tm strain when fed either a CA or AA (**Fig. 4B**). In line with previous results, WT-infected AA diet-fed mice harbored the most *S*. Tm compared to CA diet-fed mice (**Fig. 4C**). However, there was no significant difference in the pathogen burden of AA diet fed mice infected with the avirulent *S.* Tm strain fed when compared to littermates infected with the virulent WT *S.* Tm strain and fed the same diet (**Fig. 4C)**. In line with pathogen burden, we observed that AA diet-fed mice infected with WT *S.* Tm demonstrated increased ileal inflammation, assessed by histopathology, compared to CA diet-fed mice infected with either WT or avirulent *S*. Tm genotypes (**Fig. 4D)**. Surprisingly, AA diet-fed mice infected with avirulent *S*. Tm showed mild ileal inflammation (**Fig. 4D)**, suggesting that an AA diet may alter host responses to *S.* Tm in a T3SS-independent manner. Our results suggest that while the T3SSs encoded by *S*. Tm is still required to induce severe disease in the AA diet murine model as demonstrated by weight loss (**Fig. 4B**) and pathophysiology (**Fig. 4D**), T3SS activity is not needed to promote the increased *S.* Tm colonization in the ileum of AA diet-fed mice during early infection.

**Figure 4:**
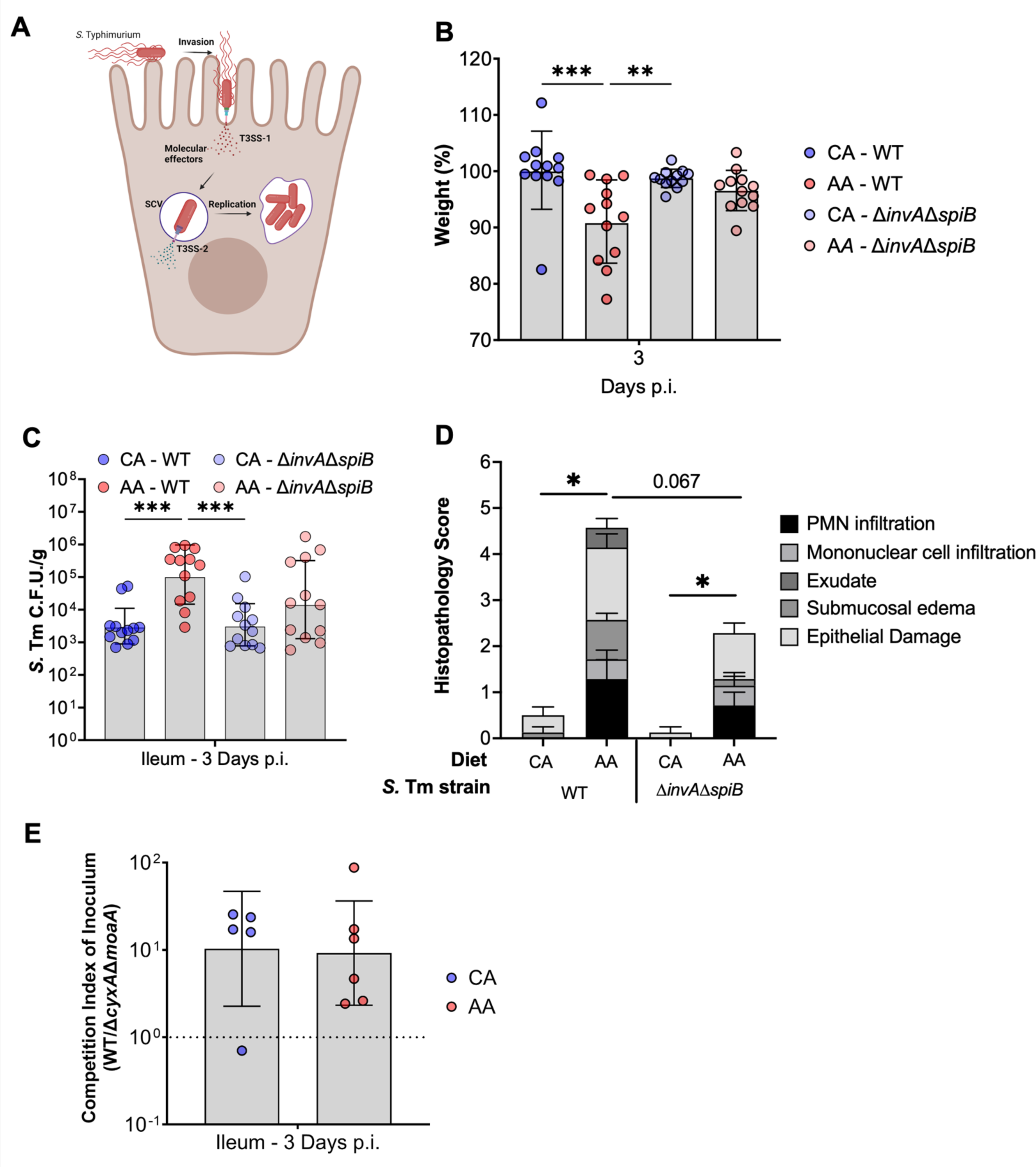
***S*. Tm does not require canonical virulence factors to colonize the ileum of AA diet fed mice.** (A) Schematic demonstrating the role of each respective T3SS used by *S*. Tm for invasion and intracellular replication in to host epithelial cells and required or the onset of gastroenteritis. (B-D) Diet-fed mice infected with either a virulent WT *S*. Tm or isogenic avirulent Δ*invA*Δ*spiB S.* Tm and carried for 3 days p.i. prior to sacrifice. (B) weight loss by day 3 p.i. and (C) luminal burden of the terminal ileum. (D) Histopathology scoring of H&E stained terminal ileum sections. Mean and SEM; Mann-Whitney. (F) CBA mice infected with a competitive ratio of WT to respiration-deficient *S*. Tm mutant, Δ*cyxA*Δ*moaA* and carried for 3 days p.i. Geometric mean and geometric SD, ANOVA or student’s t-test, n=5-12 per condition.

Models of *S.* Tm-induced large intestinal inflammation demonstrated that *S*. Tm employs anaerobic and aerobic respiration to succeed in the inflamed gut and overcome the majority of fermentative gut microbiota^47–49^. Additionally, diet alteration has been shown to impact the availability of electron acceptors in the gut, enabling pathobiont and pathogen expansion^50,51^. However, the role of anaerobic and aerobic respiration in supporting *S.* Tm colonization of the small intestine remains unknown. To interrogate if respiration was necessary for *S.* Tm ileal colonization and expansion within AA diet-fed mice, we infected mice with a 1:1 ratio of WT *S.* Tm strain and a *S.* Tm strain deficient in the ability to use inflammation-derived electron acceptors, Δ*cyxA*Δ*moaA.* Specifically, deletions in *cyxA* disable *S*. Tm from respiring low-tension oxygen, such as hypoxia, and deletions in *moaA* incapacitate the ability of *S*. Tm to respire alternative electron acceptors, such as tetrathionate, nitrate, and DSMO, due to an inability to produce a required molybdopterin cofactor required by the associated reductases^49^. Three days post-infection, we observed no differences in the fitness defects of *S.* Tm respiration-deficient isogenic mutant between diet backgrounds, suggesting an AA diet does not alter the necessity of respiration during *S.* Tm small intestinal colonization and expansion (**Fig. 4E**). Altogether, our data indicates that *S*. Tm’s expansion in the ileum of AA diet cannot be explained by increased ability of *S*. Tm to perform aerobic and anaerobic respiration.

### An AA diet significantly alters the gene expression profile of *S.* Tm in the intestinal tract *in vivo*

Considering the dispensability of the canonical virulence factors (e.g., T3SS) and no change to the requirement for aerobic or anaerobic respiration for the colonization and expansion of *S.* Tm in the ileum of AA diet-fed mice (**Fig. 4**), we turned to a more holistic approach to understand what transcriptomic changes occurred in *S*. Tm during ileal colonization of mice fed CA and AA diets. To do so, we harvested intestinal content from *S*. Tm-infected CBA/J mice fed either CA or AA diet before infection at three d.p.i. to perform *in vivo S*. Tm RNAseq. While we initially attempted to interrogate the ileum, we could not compare transcripts between diets due to the poor recovery of RNA aligning to the genome of *S.* Tm from the ileum of CA diet-fed mice, likely due to the low bacterial burden within this gut geography at this early three-day timepoint. To overcome this barrier, we used cecal content for library preparation and Illumina RNA sequencing and aligned reads to the reference genome of *S.* Tm. This was empowered by the lack of closely related species such as *E. coli* in our mice’s microbiota, as they were sourced from Jackson Labs^44^ Interestingly, we observe significant alterations in the carbohydrate and amino acid metabolism of *S.* Tm in mice fed an AA diet, with 10^7^ differentially expressed genes between CA and AA diet groups (41 genes upregulated in CA diet and 66 genes upregulated in AA diet) (**Fig. 5A**). Specifically, *S.* Tm colonizing the intestines of AA diet-fed mice demonstrated a significantly increased gene expression associated with glycosylated amino acids (the *fra* operon) (**Fig. 5B-C**), ascorbate (the *ula* operon), fructose (*frwC*) and aldose (*ego*; *yneB*) utilization (**Fig. 5C**). In addition to these degradation pathways, we also observed a significantly increased expression in amino acid biosynthesis pathways by *S.* Tm (**Fig. 5C**). Specifically, we detected a significant increase in the expression of genes associated with branched-chain amino acids biosynthesis such as valine (*ilvB*; *ilvN*), leucine (*leuB*), and isoleucine (*ilvG*; *ilvM*). *S.* Tm in the cecum content of AA-diet fed mice demonstrate significantly increased expression of the genes associated with histidine (*hisG*; *hisD*) and cysteine (*cysK*; *ybiK*) metabolism. Lastly, we observed increased expression of genes related to the central metabolism of *S.* Tm (*gldA, nadE, gltA,* and *mdh*), as well as global stressors such as acid stress or exposure to antimicrobial peptides produced by the gut epithelium or gut microbiota members such as (*uspA*; *uspC*; *uspF*; *raiA*; *zraP*; and *cspD*). Our RNAseq dataset shows significant alterations in *S.* Tm transcriptomic profile in response to AA diet, highlighting the pathogen’s adaptation to the perturbed gut microenvironment in AA diet-fed mice and identifying potential pathways that enable *S*. Tm to proliferate within the small intestine lumen.

**Figure 5:**
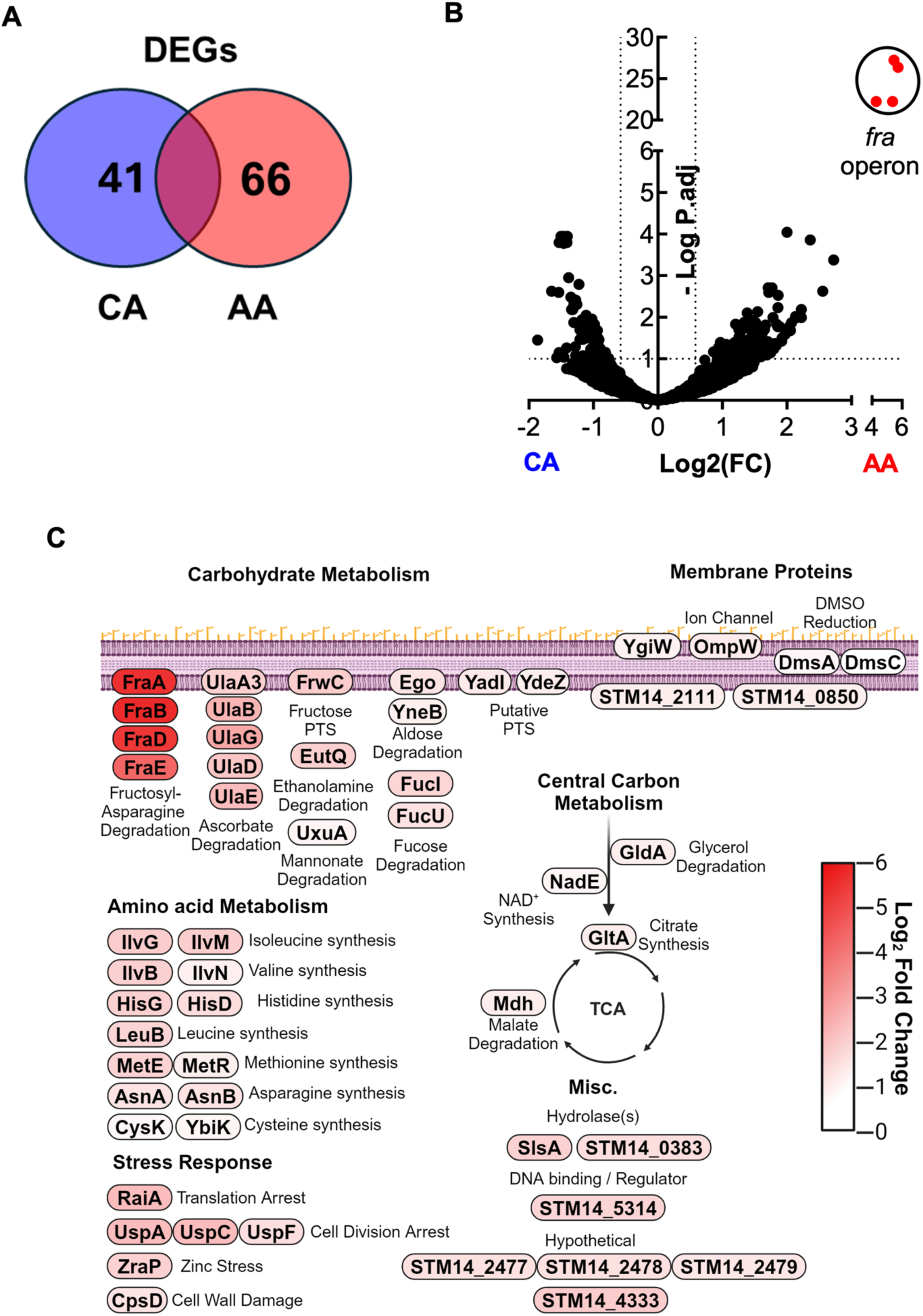
Amino acid and carbohydrate metabolism is significantly upregulated in AA diet-fed mice, 3 days p.i. (A)Venn diagram between groups of statistically significant differentially expressage genes in cecal content of S. Tm infected CBA mice fed either CA or AA diet, 3 d.p.i. (B) Volcano plot of differentially expressed *S.* Tm genes (DEGs) from the cecal content from mice switched to either a CA or AA diet two days before infection with WT *S.* Tm and carried for three days before euthanasia via DESeq2, N=3 per diet. (B) Visual schematic for the role of gene significantly upregulated in the intestine of AA diet-fed mice.

### Fructosyl-asparagine degradation significantly enables *S*. Tm ileal colonization and expansion in AA diet-fed mice

Our RNA-seq dataset identified potential pathways used by *S*. Tm to expand in the intestines of AA diet-fed mice. Since we were unable to obtain *S*. Tm transcripts from the small intestine, we performed untargeted metabolomics of the ileal content of mice fed CA or AA diets before infection as a complementary approach to our RNA-seq dataset to determine which metabolites were available to enable *S.* Tm to colonize and expand in the ileum of AA diet-fed mice during early infection. Overall, the ileal metabolic landscape of AA diet-fed mice was significantly different from that of Chow or CA diet-fed mice (**Fig. 6A, S3A-B**), with over 2600 metabolites differentially abundant in the ileal lumen of AA diet-fed mice. We detected a significant decrease in the overall abundance of bile acids and di- and tripeptides (**Fig. S3B**). AA-diet-fed mice demonstrate a significant increase in the abundance of metabolites associated with glycosylated amino acids such as Fructosyl-asparagine (F-Asn) (**Fig. 6B, S3A**). The increase in F-Asn was associated with a decrease in the abundance of its amino acid precursor, L-asparagine (**Fig. 6C**). We were specifically interested in F-Asn as our *S.* Tm *in vivo* RNAseq in intestinal content of AA diet-fed mice showed that the top DEGs were associated with the *fra* operon which enables F-Asn degradation in *S*. Tm, with *fraB* being the most highly expressed gene in our dataset (**Fig. 5B-C; 6D**). Interestingly, F-Asn is a nutrient source for *S.* Tm during intestinal inflammation^52^. However, to date, F-Asn and the *fra* operon have only been shown to be important for *S.* Tm large intestine colonization, likely due to the reduced *S*. Tm colonization of the ileum in conventional murine models of *S*. Tm intestinal disease. To confirm the importance of the *fra* operon for the colonization of *S.* Tm in the ileum of AA diet-fed mice, we constructed a mutant lacking *fraB* encoding an F-Asn deglycase. Mutants lacking *fraB* grown under conditions in which F-Asn is present as a nutrient have a reduced fitness due to the inability to detoxify intracellular 6-phosphate-fructosyl-aspartate^52^. To confirm our mutant could not use F-Asn, we cultured WT and isogenic Δ*fraB* anaerobically in LB media or in minimal media supplemented with F-Asn, *in vitro*. We note no differences in growth between WT and isogenic Δ*fraB* when cultured anaerobically in LB (**Fig. S3C**). However, when cultured in minimal media supplemented with 5mM F-Asn, Δ*fraB* demonstrated a significant growth defect compared to the WT during fermentation (**Fig. 6F**). To interrogate further the growth advantage conferred by a functional FraB *in vitro*, we repeated our kinetic growth assays comparing WT *S.* Tm to our isogenic Δ*fraB* mutant in the presence of inflammation-independent and -dependent electron acceptors such as fumarate, nitrate, tetrathionate (**Fig. 6F**), and hypoxia (**Fig. 6E**). We found that deletion of *fraB* conferred a growth defect in all conditions where F-Asn was the major nutrient source for *S.* Tm independent of fermentation or respiration. To determine the fitness cost of mutants lacking *fraB in* vivo in the different segments of the intestinal tract, we repeated our diet model by infecting mice with a 1:1 ratio of WT and our *fraB* deficient strain (Δ*fraB*). Interestingly, at three days post-infection, only mice pre-fed our AA diet demonstrated a ten-fold defect of the Δ*fraB* mutant compared to the CA diet-fed mice in the ileum (**Fig. 6G-H**). Similar results were observed in the cecum and colon of AA diet-fed mice (**Fig. 6G**). Taken together, these data suggest that F-Asn is an important nutrient source for the expansion of *S*. Tm in the ileum of AA diet-fed mice during early infection and that the inability of the Δ*fraB* mutant to colonize the ileum may impact its fitness in the large intestine.

**Figure 6:**
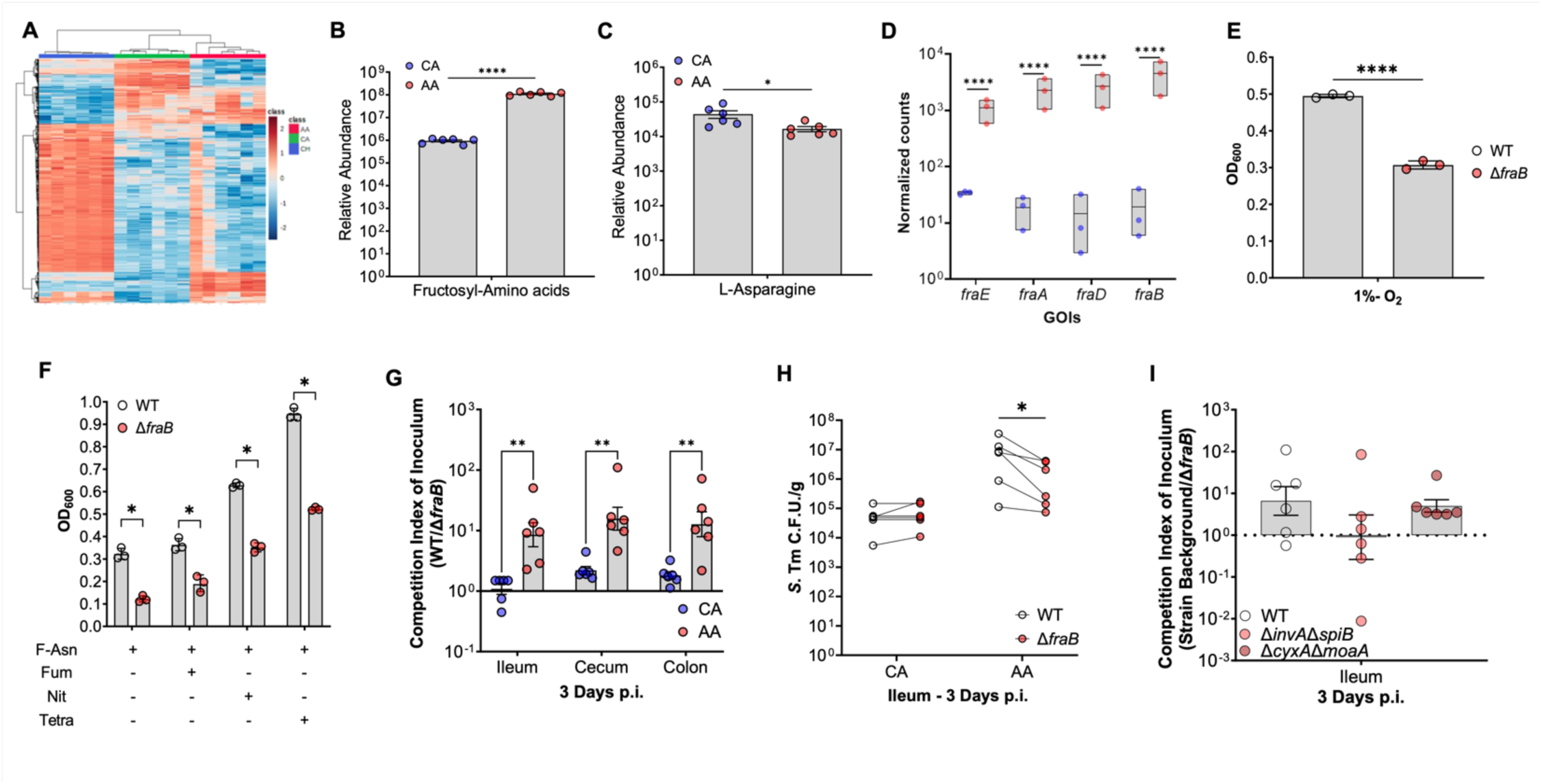
AA diet increases F-Asn bioavailability in the terminal ileum before inflammation onset, enabling expansion of *S*. Tm. (A) Heatmap of metabolic profile of ileal content of uninfected mice maintained on Chow or switched to either a CA or AA-based diet for two days before euthanasia, N=6. (B-C) Relative abundance of (B) fructosylated amino acids and (C) L-asparagine from the terminal ileum contents of CA or AA diet-fed uninfected mice. N=6, PERMANOVA. (D) Transcript abundance for RNAseq from CA or AA diet-fed mice contents three days p.i. associated with the degradation and utilization of fructosyl-asparagine, the *fra* operon. N=3. To interrogate the ability of *S.* Tm to grow in the presence of fructosyl-asparagine, we cultured WT and an isogenic mutant lacking the F-Asn deglycase encoded by *fraB* (E) under hypoxia like that observed in the small intestine and (F) anaerobically with and without the addition of inflammation derived electron acceptors (fumarate, nitrate, or tetrathionate; 40mM) associated with large intestine expansion. Optical density at 600nm was measured hourly in the respective conditions. Data represents the average of three biological replications (each the mean of three technical replicates). Student’s t-test, P=*, <0.05; **, <0.01; ***, <0.001; ****, <0.0001. To interrogate the fitness cost of *fraB, in vivo,* we infected CBA mice switched to either a CA or AA diet two days before infection with a competitive ratio of WT: Δ*fraB* marked via differential antibiotic resistance and carried for three days post-infection. (G) Competition index of inoculum of WT to *fraB* from the terminal ileum, cecum, and colon 3 d.p.i.; N=6 per diet, Multiple t-test, P=**, <0.01. (H) CFU enumeration of competitive infection, comparing WT and Δ*fraB* isogenic *S.* Tm strains in CA and AA diet fed mice. Paired Multiple t-test, P=*, <0.05. (I) Competition index of inoculum in WT, avirulent (Δ*invA*Δ*spiB*), and respiration-deficient (Δ*cyxA*Δ*moaA*) backgrounds. N=6 per strain background. ANOVA.

In addition to interrogating the necessity of the *fra* operon in the ability of *S*. Tm to colonize the ileum of AA-diet fed mice, we constructed mutants in the other top four DEGs and pathways upregulated in AA-diet fed mice within our *in vivo S.* Tm RNAseq dataset (**Fig. 5**). To do so, we generated *S.* Tm isogenic mutants in the following genes: the ribosomal stall protein RaiA encoded by *yfiA*; the zinc-stress protection protein ZraP encoded by *zraP*; part of the L-ascorbate-specific phospho-transfer system (PTS) EIIC component SgaB encoded by *ulaA_3;* and lastly the fructose-specific PTS EIIC component FrwC encoded by *frwC.* Like *fraB*-deficient mutants, deletions in *yfiA, zraP*, *ulaA_3,* or *frwC* did not confer an overall *S.* Tm growth defect when cultured in LB anaerobically, *in vitro* (**Fig. S3C**). To assess the necessity of each of these genes or metabolic pathways within the ileum of AA-diet-fed CBA mice, we infected CA- and AA-diet-fed mice with a 1:1 competitive ratio of WT and either a Δ*yfiA*, Δ*zraP*, Δ*ulaA_3*, or Δ*frwC.* Interestingly, we observed that similarly to deletions in *fraB*, deletions in *yfiA* resulted in a significant fitness defect only in the ileum of AA-diet-fed mice. No significant fitness defects were observed in mutants deficient in *zraP, ulaA_3,* or *frwC* (**Fig.S3D**). While our data indicates that WT *S.* Tm can outcompete the isogenic Δ*yfiA* in the ileum of AA diet-fed mice, we find significant fitness advantage of WT *S.* Tm over an isogenic *ΔyfiA* diets in the large intestine (cecum and colon) of both CA and AA diet fed animals, suggesting the requirement of *yfiA* to be diet independent in the large intestines (**Fig. S3E**). Taken together, these data suggest that the *fra* operon may be the most influential in supporting *S*. Tm colonization in the ileum of AA diet-fed mice during early infection.

Previous work by Ali *et al.* demonstrated using GF and streptomycin-treated conventional C57Bl/6 mice that *S*. Tm requires the action of its two T3SS as well as the ability to respire inflammation-derived tetrathionate to take advantage of F-Asn in the gut. However, F-Asn enabled *S.* Tm growth in the absence of electron acceptors *in vitro*, thus suggesting that F-Asn could support *S.* Tm growth before the onset of inflammation. Given the dispensability of the T3SS and respiration in the ability of *S.* Tm to colonize and expand within the ileum of AA diet fed mice in **Fig. 4**, we aimed to interrogate the reliance of *S.* Tm on its primary virulence program and respiration in the use of F-Asn in the ileum. To do so, we took advantage of the avirulent background of *S.* Tm, Δ*invA*Δ*spiB* which is unable to cause severe disease or to gain access to inflammation-derived electron acceptors^41^; and constructed an isogenic Δ*fraB* deletion strain in this background (Δ*invA*Δ*spiB*Δ*fraB).* We subsequently repeated our murine infection in AA-diet-fed mice, competing S. Tm Δ*invA*Δ*spiB* with its isogenic Δ*invA*Δ*spiB*Δ*fraB* mutant. We note no significant differences in the defect of *fraB* deficient strain between the WT and avirulent backgrounds in line with our previous results that *S.* Tm does not require its T3SS to expand in the ileum of AA diet-fed mice. Additionally, to target both aerobic and anaerobic respiration, we made a *fraB* deletion in a *S.* Tm background lacking the low-affinity oxygen reductase, Δ*cyxA,* and the rate-limiting step in the synthesis of molybdopterin cofactors, Δ*moaA*, which is required for utilization of alternative electron acceptors, thus generating the strain Δ*cyxA*Δ*moaA*Δ*fraB.* We repeated our competitive infections with the aim of understanding if aerobic or anaerobic respiration was required for the *S*. Tm’s ability to use F-Asn in the ileum of our AA diet fed mice. No significant differences were observed the ability of respiration-deficient *S*. Tm background strains to use F-Asn when compared to WT *S.* Tm (**Fig. 6I**). Taken together our data demonstrates that an AA diet increases the availability of a microbiota-sequestered nutrient source, F-Asn which supports *S.* Tm expansion in the ileum before the onset of inflammation in a respiration-independent manner.

## DISCUSSION

For many years, the study of *Salmonella enterica* serovar Typhimurium has illuminated many factors that impart an advantage of enteric pathogens over the resident collection of gut commensals making up the gut microbiota. However, until recently, the dynamics between *S*. Tm and the small intestine microbiota remained to be fully understood due to critical limitations in methodologies and effective animal models^17,28^. Herein we describe a novel mouse model of diet-dependent manipulation of the gut microbiota to study the pathogenesis of the enteric pathogen *Salmonella enterica* serovar Typhimurium during intestinal infection within the context of the small intestine. Feeding genetically resistant mice (CBA/J) a novel L-amino acid diet remarkably enabled *S*. Tm colonization and expansion in the ileum resembling features of *S.* Tm infection seen in humans. Using our model, in combination with multi-omics approaches, we discovered new pathways used by *S*. Tm to colonize the lumen of the small intestine. Further, we found that conventional virulence factors (T3SS and respiration) important for *S.* Tm success in the large intestine were not required for *S*. Tm to colonize the small intestine of AA diet-fed mice. Thus, these findings necessitate further research into small intestine-specific factors that influence *S*. Tm success in this distinct geography from that of the large intestine, such as the role of fimbriae. We confirm previous findings from studies focused on metabolic mechanisms of *S*. Tm intestinal colonization of the large intestine that apply to the small intestine. Specifically, we validate the use of our model by demonstrating that, similarly to the large intestine, *S.* Tm benefits from the use of Fructosyl-asparagine (F-Asn) to expand in the small intestine.

Interestingly, work by Sabag-Daigle demonstrates that the capacity to degrade F-Asn is found in multiple bacterial phyla, including Actinobacteria, Firmicutes, and other Proteobacteria (i.e., *Citrobacter* spp., *Klebsiella* spp.,) ^52,53^. While reconstitution of AA diet mice with a complex microbiota was sufficient to alter the *S.* Tm colonization and expansion of the ileum in our model, pre-colonization of mice with commensal Proteobacteria did not significantly alter the ability of *S*. Tm to expand in the ileum. Work by Ali *et al.* has shown that the *fra* operon is not highly conserved and represents horizontal gene acquisition for some pathogenic strains^54^. Thus, this suggests that one reason commensal *Enterobacteriaceae* were not able to compete with *S*. Tm in the ileum was likely due to the lack of the necessary *fra* operon within the chosen representatives. Using a commensal representative that encodes this operon will provide much-needed research to understand if F-Asn is the only nutrient made available to *S.* Tm in the ileum of AA diet-fed mice.

In addition to the significant differential expression of the *fra*, we observed several other operons expressed by *S.* Tm in the cecum of AA diet mice. Specifically, the *ula* operon associated with the import and degradation of ascorbic acid were significantly expressed in the intestinal content of AA diet-fed mice. However, unlike the observation of the necessity of *fra*, we observed no fitness defects of a mutant unable to transport ascorbic acid *in vivo* at our early day three timepoint, despite confirming its defect *in vitro (data not shown).* Previous reports by Teafatiller *et al*. suggest that mice infected with *S.* Tm accumulate ascorbic acid in the intestinal lumen due to inflammation-dependent decreased expression of the transporter SVCT1^55^. Many inflammation-dependent phenotypes do not demonstrate a significant difference before the onset of inflammation. Thus, further studies are required to understand the contribution of ascorbic acid degradation on *S*. Tm pathogenesis. In addition to the *ula* operon, we observed a significant increase in the expression of several amino acid biosynthesis pathways within *S*. Tm. While work from Gabrielle-Nunez et al. has demonstrated that pathogens utilize amino acid biosynthesis to subvert amino acid sequestration by the gut microbiota^56^. Which amino acids and how *S*. Tm overcomes this effective mechanism of imparting nutrient scarcity remains to be fully investigated. Use of diet-induced S. Tm ileitis may help shed light on the influences of each amino acids and other nutrients and metabolites in the context small intestine *S.* Tm colonization as well as dissect the contribution of microbiota constituents to ileal-specific mechanisms of microbiota-mediated colonization resistance to *S*. Tm infection.

## Acknowledgments

N.G.S. was supported by NIH T32 Training Grant (T32ES007028-46) and HHMI Gilliam Fellowship (GT15104). T.M.R. was supported by ACS Postdoctoral Fellowship (PF-24-1247878-01-CCB). K.J. was supported by NIH T32 Training Grant (T32GM13779). M.X.B is a HHMI Freeman Hrabowski Scholar. Work in M.X.B.’s laboratory was supported by the NIH (R01DK131104 and 1R01AI168302), The Pew Charitable Trusts (2022-A-19568) and the Burroughs Wellcome Fund (022792).

## Author contributions

Conceptualization, N.G.S., and M.X.B.; Methodology, N.G.S., and M.X.B.; Investigation, N.G.S., M.B., C. B., P.C., A.C.S., S.C., S. S., S.J., T. M. R.; K. J., J.M., D. B.J; Writing – Original Draft, N.G.S. and M.X.B.; Writing – Review & Editing, N.G.S., M.B., C. B., P.C., A.C.S., S.C., S. S., S.J., T. M. R.; K. J., J.M., D. B.J; M.X.B.; Funding Acquisition, M.X.B.; Resources, M.X.B.; Supervision, M.X.B.

## STAR METHODS

### RESOURCES AVAILABILITY

#### Lead contact

Further information and requests for resources and reagents should be directed to and will be fulfilled by the lead contact, Mariana X. Byndloss.

#### Materials availability

All unique reagents generated in this study are available from the lead contact without restriction.

#### Data and code availability

Data reported in this paper will be shared by the lead contact upon request. This paper does not report original code. Any additional information required to reanalyze the data reported in this paper is available from the lead contact upon request. The untargeted metabolomics data is available at the NIH Common Fund’s National Metabolomics Data Repository (NMDR) website, the Metabolomics Workbench, https://www.metabolomicsworkbench.org, under the assigned Study ID ST003796. The data can be accessed directly via its Project DOI: http://dx.doi.org/10.21228/M8582Z. Ileum bulk RNAseq, intestinal content RNAseq, as well as 16S rRNA raw reads available associated with BioProject PRJNA1241294.

### EXPERIMENTAL MODEL AND SUBJECT DETAILS

#### Mouse husbandry

All animal experiments in this study were approved by the Institutional Animal Care and Use Committee at the Vanderbilt University Medical Center. Female CBA mice were obtained from the Jackson Laboratory. Germ-Free Swiss Webster (SW) mice were bred in-house and maintained in specific pathogen-free facilities at Vanderbilt University Medical Center. A subset of germ-free mice was colonized with a defined microbiota to obtain the SW mice used in this study. The age at the beginning of the experiment was 6-7 weeks old for all CBA/J and SW mice. Conventional mice were housed in individually ventilated cages with ad *libitum* access to water and chow (Pico Labs Inc. 5L0D Diet). Germ-free SW mice were housed in sterile positive pressure cages (Allentown) with ad *libitum* access to sterile water and chow.

Unless indicated otherwise in the figure legend, male and female mice were analyzed for the germ-free SW mice, and no significant sex-specific differences were noted. Both sexes were equally represented in each experimental group. Animals were randomly assigned into cages and treatment groups 3 days prior to experimentation. Unless stated otherwise, a minimum of 5 mice were used based on variability observed in previous experiments. All mice were monitored daily, and cage bedding changed every two weeks. At the end of the experiments, mice were humanely euthanized using carbon dioxide inhalation. Animals that had to be euthanized for humane reasons prior to reaching the predetermined time point were excluded from the analysis.

#### Bacterial Strains

*S*. Typhimurium IR715, *E. coli* Mt1B1, *K. pneumoniae* strains were routinely grown aerobically at 37°C in LB broth (10 g/L tryptone, 5 g/L yeast extract, 10 g/L sodium chloride) or on LB agar plates (10 g/L tryptone, 5 g/L yeast extract, 10 g/L sodium chloride, 15 g/L agar). *P. mirabilis* was routinely grown aerobically at 37°C on MacConkey (BD Difco) agar or in LB broth. When appropriate, agar plates and media were supplemented with 30 mg/mL chloramphenicol (Cm), 100 mg/mL carbenicillin (Carb), 100 mg/mL kanamycin (Km), or 50 mg/mL nalidixic acid (Nal).

*<u>Defined Microbiota</u>*: *Muribaculum intestinale*, *Bacteroides caecimuris*, *Clostridium clostridioforme*, *Clostridium innocuum*, *Clostridioides mangenotti*, *Clostridium cochlearium* and *Clostridium sporogenes, Paraclostridium bifermentans, Intestinimonas butyriciproducens, Blautia coccoides, Akkermansia muciniphila* were cultured on Columbia agar (BD Difco) supplemented with 5% defibrinated sheep blood or Akkermansia Anerobic Media (18.5g/L Brain-Heart Infusion, 5g/L Yeast Extract, 15g/L Trypsin-Soy Broth, 2.5g/L Potassium monophosphate, 0.5g/L Glucose, 0.1g/L Sodium Bicarbonate, 0.5g/L Cysteine-HCl, 3% Heat-inactivated Fetal Cafl/Bovine Serum, 5mg/L Menadione, 1mg/L Hemin). *Lactobacillus reuteri* and *Bifidobacterium longum* subsp. *animalis* were cultured on MRS agar or broth. *Staphylococcus xylosus,* and *Streptococcus danieliae* Tryptic Soy agar or broth in an anaerobic chamber (85% nitrogen, 10% hydrogen, 5% carbon dioxide, Coy Lab Products). All media was place into anaerobic chamber for at least 48hrs to pre-reduce prior to culture of any organism.

### METHODS DETAILS

#### Construction of plasmids and bacterial strains

*S.* Tm mutant construction was done via lambda red recombination as previously described^57^. In brief, primers were designed to be 40bps homologous regions upstream and downstream of the gene of interest and used to amplify the Kan^R^ cassette in pKD13. Kan cassette was amplified, and PCR product DpnI was treated and purified before electroporation into S. Tm IR715 harboring the helper plasmid pKD46 grown in the presence of 0.1M L-arabinose. The transformation was plated on LB agar supplemented with Kanamycin and incubated overnight at 37°C and colonies were checked via PCR. Following insertion confirmation, insertion was pushed into a clean background as previously described^58,59^ before transformation with pCP20 to remove the Kan^R^ cassette.

Alternatively for the deletion of *zraP* and *yfiA*, suicide vectors were constructed by digesting pGP704 with restriction enzyme EcoRV-HF, amplifying the 500bp up- and downstream of target gene ORF, and ligated using HiFi assembly. Assembly was then transformed into electrocompetent DH5α-λ*pir* and clones assayed via PCR for 1kb insert. Vectors then confirmed via sequencing, before transformation into conjugative S17-λ*pir* and mated with WT *S*. Tm IR715. Merodiploid *S*. Tm selected for using a combination of Nal and Kan supplementation in LB agar and confirmed via PCR. Following sucrose selection, mutants then confirmed via PCR for KO.

#### Growth Kinetics

Bacteria were struck out onto fresh selective agar plates and incubated overnight. Single colonies were taken and cultured in LB broth supplemented with appropriate antibiotics in biological triplicate for three independent experiments. Overnight cultures were then back diluted 1:100 for 1-2 hours in 5mL of fresh LB broth supplemented with antibiotics until the bacteria reached the mid-log phase. Cultures were then washed twice and normalized to an OD_600_=1. Cultures were then inoculated into a final volume of 200uL per well of pre-reduced media to a final OD_600_=0.01, shaking continuously, incubated at 37°, and OD_600_ measured every hour. When necessary, carbon sources (Glucose or F-Asn) were supplemented at 5mM, while electron acceptors (Fumarate, Nitrate, and Tetrathionate) were supplemented at 40mM.

#### Animal Experiments

Animal models of *S.* Typhimurium colitis

##### CBA mouse model

CBA mice were infected with either 0.1 mL of LB broth (mock-infected) or *S*. Typhimurium IR715 strains in LB broth. Groups (n=4-5) of 6-7 weeks CBA mice were intragastrically infected with 1 x 10^7^ CFU for single strain and competitive infection experiments. Fecal samples were collected on days 1, 3, 5, and 7 days after infection for bacterial plating. Three or seven days after infection, samples for histopathology, terminal ileal luminal content, colonic luminal content, and cecal content for *S.* Tm enumeration. *S.* Tm numbers were determined by plating serial ten-fold dilutions onto LB agar containing the appropriate antibiotics. For the aspartate-free diet experiments, Groups (n=5) of 6-7 weeks old CBA mice were fed control diet A20073101 (AA Diet; Research Diets) or D22011804 (Casein Diet; Research Diets) beginning 48h before *S.* Tm infection and were kept on the diets throughout the experiment. The remaining experiment procedures were performed as described above.

##### Germ-Free Swiss Webster (SW) mice

Groups (n=5) of 6-7 weeks old SW germ-free mice were intragastrically inoculated with 1 x 10^9^ CFU containing a an equal mixture of *Muribaculum intestinale*, *Bacteroides caecimuris*, *Lactobacillus reuteri*, *Bifidobacterium longum* subsp. *animalis*, *Clostridium clostridioforme*, *Clostridium innocuum*, *Clostridioides mangenotti*, *Clostridium cochlearium* and *Clostridium sporogenes, Paraclostridium bifermentans, Intestinimonas butyriciproducens, Blautia coccoides, Akkermansia muciniphila, Staphylococcus xylosus,* and *Streptococcus danieliae.* After 5 days, pre-colonized and germ-free control animals were switched to double gamma-irradiated Chow, CA, or AA diets for 2 days prior to infection with 1 x 10^7^ *Salmonella enterica* serovar Typhimurium IR715. Three days after infection, samples for histopathology, colonic luminal content and cecal content for *S.* Tm enumeration. *S.* Tm numbers were determined by plating serial ten-fold dilutions onto LB agar containing the appropriate antibiotics.

#### Animal model of *S*. Typhimurium intraperitoneal infection

Two days before infections, cultures were struck out on to fresh agar plates supplemented with appropriate antibiotics, then a single colony used for an overnight culture in 3mL of LB liquid broth. The day of the infection, cultures were harvested via centrifugation before being washed twice with sterile 1X PBS and normalized to the desired OD_600_, based on route of infection. After infection, inoculum bacterial density enumerated via serial dilution on agar plates supplemented with appropriate antibiotics for CFU counting.

#### Histopathology scoring

Ileal tissue was fixed in 10% phosphate-buffered formalin for 48 h and embedded in paraffin. Sections (5 µm) were stained with hematoxylin and eosin. Stained sections were blinded and evaluated by a veterinary pathologist according to the criteria listed in **Table S7**.

#### Bacterial 16S rRNA sequencing

Small intestinal content was collected at sacrifice, flash-frozen in liquid nitrogen, and stored at −80°C before processing (n=9, 3 per diet). DNA was then extracted from the samples using the DNeasy PowerSoil Kit (Qiagen #47016) according to the manufacturer’s instructions. Sequencing libraries were prepared by the University of California, San Diego (UCSD) Microbiome Core using protocols and primers published on the Earth Microbiome Project website and sequencing performed on Illumina NovaSeq 6000 according to the manufacturer’s instructions. Other samples were sequenced by Seqmatic on Illumina MiSeq. Paired end sequences were imported into QIIME2 (version QIIME2-2023.2). Reads were trimmed to exclude regions that fell below a quality score of 25. Features tables for each data set were constructed and then merged following denoising. Data was rarefied based on minimum read quantity. Phylogenetic trees were then constructed, and taxonomy was assigned based on the Silva genes classifier. Pseudocounts added and ANCOM was performed to identify alterations in gut microbiota composition based on diet or administration of low-dose penicillin. Feature tables constructed for taxonomic levels 5(genera), 6(family), and 2(phylum). Pseudocounts added to each of the collapsed feature tables and ANCOMs performed at for each experimental variable as previously performed. Distance matrices (jaccard, bray-curtis, weighted, and unweighted) were exported from QIIME2 for further processing in R-studio. Alpha diversity metrics (Shannon, Faith_pd, and Observed features) were extracted from QIIME2 for downstream applications. PERMANOVAs performed for each distance matrix, as well as the creation of non-metric multidimensional scaling (NMDS) plots based on each distance matrix.

#### Untargeted metabolomics on small intestinal content

Small intestinal content from mice was collected following a diet switch and maintenance for two days without infection (n=18, 6 per diet) prior to sacrifice, snap-frozen, and stored at −80°C. Small intestinal content from mice was collected following a diet switch and maintenance for two days without infection (n=18, 6 per diet) prior to sacrifice, snap-frozen, and stored at −80°C. Individual frozen small intestinal content was lysed in 1000 μl ice-cold lysis buffer (1:1:2, v:v:v, acetonitrile: methanol (MeOH): ammonium bicarbonate 0.1M -pH 8.0) and sonicated individually using a probe tip sonicator at 50% power (10 pulses). Isotopically labeled standards (phenylalanine and biotin) were added to each sample to determine sample process variability as previously described)^60–63^. Homogenized samples were normalized by weight to the smallest amount of intestinal content such that each sample contained an equal amount of intestinal content into a 200 µl final volume of lysate. Proteins were precipitated from individual samples by adding 800 µL of ice-cold MeOH followed by overnight incubation at −80°C. Precipitated proteins were pelleted by centrifugation (10k rpm, 10 min) and extracts were dried down *in vacuo* and stored at −80°C.

Extracted samples were reconstituted in 100 µL H_2_O, and 100 µL MeOH with vortex mixing after each addition. Samples were incubated at room temperature for 10 min followed by liquid-liquid extraction. For liquid-liquid extraction (LLE), 800 µL MTBE was added with vortex mixing for 30 sec followed by incubation on ice for 10 min and centrifugation at 15k rpm for 15 min at 4°C. The lower (hydrophilic) fraction was transferred into a new Eppendorf tube, dried *in vacuo*, and stored at −80 °C until further use.

Prior to mass spectrometry analysis, individual hydrophilic extracts were reconstituted in 100 µL acetonitrile/water (80:20, v/v) containing isotopically labeled standards, tryptophan, inosine, valine, and carnitine, and centrifuged for 5 min at 10,000 rpm to remove insoluble material. A pooled quality control (QC) sample was prepared by pooling equal volumes of individual samples following reconstitution. The QC sample allowed for column conditioning (eight injections), retention time alignment, and assessment of mass spectrometry instrument reproducibility throughout the sample set.

LC-MS and LC-MS/MS analyses were performed on a high-resolution Q-Exactive HF hybrid quadrupole-Orbitrap mass spectrometer (Thermo Fisher Scientific, Bremen, Germany) equipped with a Vanquish UHPLC binary system (Thermo Fisher Scientific, Bremen, Germany). Extracts (5 μL injection volume) were separated on an ACQUITY UPLC BEH Amide HILIC 1.7 μm, 2.1 × 100 mm column (Waters Corporation, Milford, MA) held at 30°C using the LC method as previously described. Full MS analyses were acquired over the mass-to-charge ratio (*m/z*) range of 70 - 1,050 in positive ion mode. The full mass scan was acquired at 120K resolution with a scan rate of 3.5 Hz and an automatic gain control (AGC) target of 1×10^6^. Tandem MS spectra were collected at 15 K resolution, AGC target of 2×10^5^ ions, and maximum ion injection time of 100 ms. Data analysis was ^64–67^performed as previously described^68^. Briefly, spectral features (retention time, m/z pairs) were extracted from the data using Progenesis QI v.3.0 (Non-linear Dynamics, Waters Corporation, Milford, MA). Both MS and MS/MS sample runs were aligned against a pooled QC sample reference run, and features were deconvoluted. Data were normalized to all compounds, and species with >25% coefficient of variation (CV) in the pooled QC samples were removed from analysis. Global assessments were performed in Metaboanalyst 5.0^69^. Accurate mass measurements (< 5 ppm error), isotope distribution similarity, fragmentation spectrum matching against the Human Metabolome Database (HMDB)^70^ and both fragmentation spectrum and retention time matching to an in-house library of reference standards were used to determine tentative, putative, and validated (Level 1-3) annotations^70^.

#### Synthesis of fructosyl-asparagine

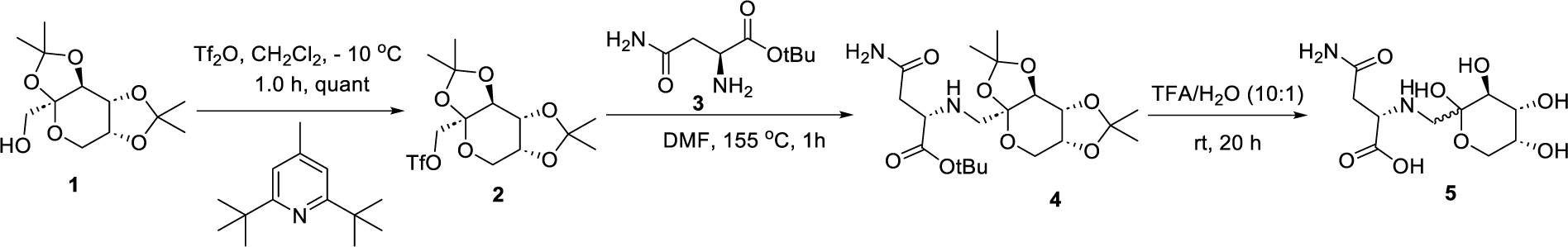

**((3a*S*,5a*R*,8a*R*,8b*S*)-2,2,7,7-tetramethyltetrahydro-3a*H*-bis([1,3]dioxolo)[4,5-b:4’,5’-d]pyran-3a-yl)methyl trifluoromethanesulfonate (2)**

At −10 °C, trifluoromethanesulfonic acid anhydride (1.35 mL, 11.0 mmol) was added dropwise under an atmosphere of nitrogen to a solution of 2,6-di-*tert*-butyl-4-methylpyridine (1.3 mL, 7.7 mmol) in dry dichloromethane (30 mL). To this mixture was added dropwise a solution of ((3aS,5aR,8aR,8bS)-2,2,7,7-tetramethyltetrahydro-3aH-bis([1,3]dioxolo)[4,5-b:4’,5’-d]pyran-3a-yl)methanol (**1**, 1.0 g, 3.8 mmol) in dichloromethane (10 mL) while stirring. The reaction mixture was stirred for 1 h at −10 °C before adding ice-cooled water. The solution was extracted with dichloromethane (3 x 30 mL). The combined organic extracts were dried over Na_2_SO_4_, filtered, and the solvent was evaporated. The chromatography of the crude product on silica gel (hexane/ethyl acetate, 4:1) and evaporation of the solvent from the appropriate fractions gave **2** (1.5 g, quantitative) as a yellow syrup. ^1^H NMR (400 MHz, CDCl_3_) 4.66 (dd, *J* = 2.7, 7.9 Hz, 1H), 4.54 (d, *J* = 10.5 Hz, 1H), 4.43 (d, *J* = 10.3 Hz, 1H), 4.33 (d, *J* = 2.7 Hz, 1H), 4.27 (dd, *J* = 1.2, 7.9 Hz, 1H), 3.95 (dd, *J* = 1.8 Hz, 13.0 Hz, 1H), 3.81 (dd, *J* = 0.4, 12.9 Hz, 1H), 1.58 (s, 3H), 1.49 (s, 3H), 1.42 (s, 3H), 1.37 (s, 3H).

#### *tert*-butyl (((3a*S*,8b*S*)-2,2,7,7-tetramethyltetrahydro-3a*H*-bis([1,3]dioxolo)[4,5-b:4’,5’-*d*]pyran-3a-yl)methyl)-L-asparaginate (4)

The triflate derivative **2** (650.0 mg, 1.65 mmol) was dissolved in anhydrous DMF (20 mL), and *tert*-butyl L-asparaginate (**3**, 623.0 mg, 3.31 mmol) was added to the solution. The mixture was heated at 155 °C for 1 h, The reaction was quenched by adding ice-water (50 mL), and the aqueous layer was extracted with Ethyl acetate (3 x 50 mL). The combined organic layer was washed with ice-water (3 x 30 mL), and the solvent was evaporated under reduced pressure. The column chromatography of the crude product on silica gel (MeOH/DCM 3:97) and evaporation of the solvent from the appropriate fractions yielded **4** (270 mg, 38%). ^1^H-NMR (400 MHz, CDCl_3_) 7.91 (s, 1H), 5.55 (s, 1H), 4.48 (d, *J* = 8.1 Hz, 1H), 4.15 (d, *J* = 12.5 Hz, 2H), 3.79 (d, *J* = 13.1 Hz, 1H), 3.68 (d, *J* = 12.5 Hz, 1H), 3.40 (d, *J* = 8.9 Hz, 1H), 2.97 (d, *J* = 13.4 Hz, 1H), 2.67 (d, *J* = 11.7 Hz, 1H), 2.47 (dd, *J* = 3.3, 15.7 Hz, 1H), 2.33 (dd, *J* = 9.5, 16.2 Hz, 1H), 1.45 (s, 3H), 1.40 (s, 3H), 1.38 (s, 9H), 1.30 (s, 3H), 1.26 (s, 3H). LCMS: Rt = 0.585 min, M+H =431.1, 1 min method.

#### (((3*S*,4*R*,5*R*)-2,3,4,5-tetrahydroxytetrahydro-2*H*-pyran-2-yl)methyl)-L-asparagine (5)

The compound **4** (350.0 mg, 0.81 mmol) was dissolved in trifluoroacetic acid/water (9:1, v/v, 8.9 mL) and stirred for 16 h at room temperature. The solvent was evaporated in vacuo. The crude was dissolved in 5.0 mL MeOH (minimum amount), added diethyl ether (10 mL) to precipitate out the product, filtered off and washed with ether. The solid was kept in high vacuum for 16 h to provide **5** (142.0 mg, 59%, mixture of anomeric isomers) as a white solid. ^1^H-NMR (400 MHz, D_2_O) 4.21-4.01 (m, 1H), 3.99-3.89 (m, 1H), 3.85-3.57 (m, 3H), 3.53-3.24 (m, 3H), 3.04-2.87 (m, 2H). LCMS: Rt = 0.083 min, M+H =295.1, 1 min low mass method.

#### Host bulk RNA sequencing

Ileal tissue was collected at sacrifice and fresh frozen in liquid nitrogen before storage at −80°C. RNA was isolated using a combination of TRIzol and Purelink RNA extraction kit per manufacturer’s instructions. RNA was treated with TURBO DNase before submission to VANTAGE sequencing. Following sequencing, STAR was used to align sequencing to the reference genome of CBA mice, and then subsequently, used for FeatureCounts to enumerate the transcripts identified for each sample. DESeq2 was then utilized to compare between groups and identify differentially expressed genes.

#### Gene expression from host tissue

Ileal tissue was isolated at necropsy at the represented timepoints and snap frozen in liquid nitrogen before storage at −80°C prior to RNA isolation. Using a combination of TRIzol and lysing matrix B tubes (MPBio), tissue homogenized and RNA isolate using the PureLink RNA isolation kit according to manufacturer’s instructions. cDNA synthesized form 1µg of RNA, following DNase treatment using iScript gDNA clear cDNA synthesis kit, and then subsequently run using iQ SYBR green Supermix for interrogation of host genes.

#### Intestinal Content pH measurement

Ileal luminal content harvested from mice fed either a CA or AA diet for seven days prior to euthanasia, and resuspended at 100mg/mL in ddH2O prior to measurement of pH via pH strips (Fisherbrand^TM^).

#### Bacterial bulk RNA sequencing

Intestinal content was collected at sacrifice three days after infection, flash-frozen in liquid nitrogen, and stored at −80°C. RNA was extracted using the RNeasy Plus Mini Kit and used for library preparation using the Illumina Total RNA Stranded Ligation kit with Ribo-Zero Plus regent for depletion of both host and bacterial (gram negative and gram positive) ribosomal RNA. Quality assessment and quantification of RNA preparations and libraries were performed using an Agilent 4200 TapeStation and Qubit 3, respectively. Sequencing was carried out on an Illumina NextSeq 2000 instrument using a 300 cycle kit to generate paired-end 150 bp sequence reads.

### KEY RESOURCES TABLE

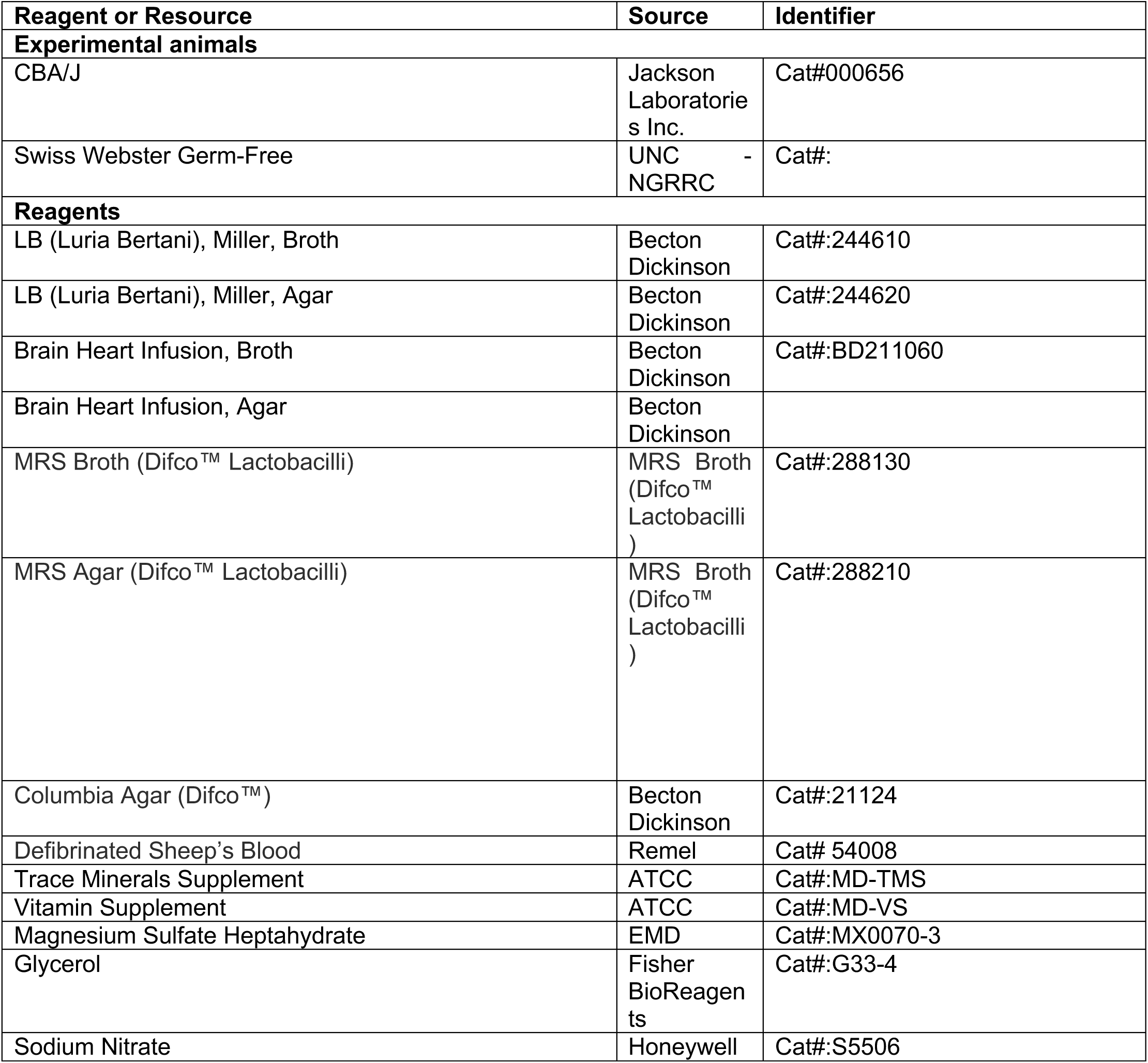
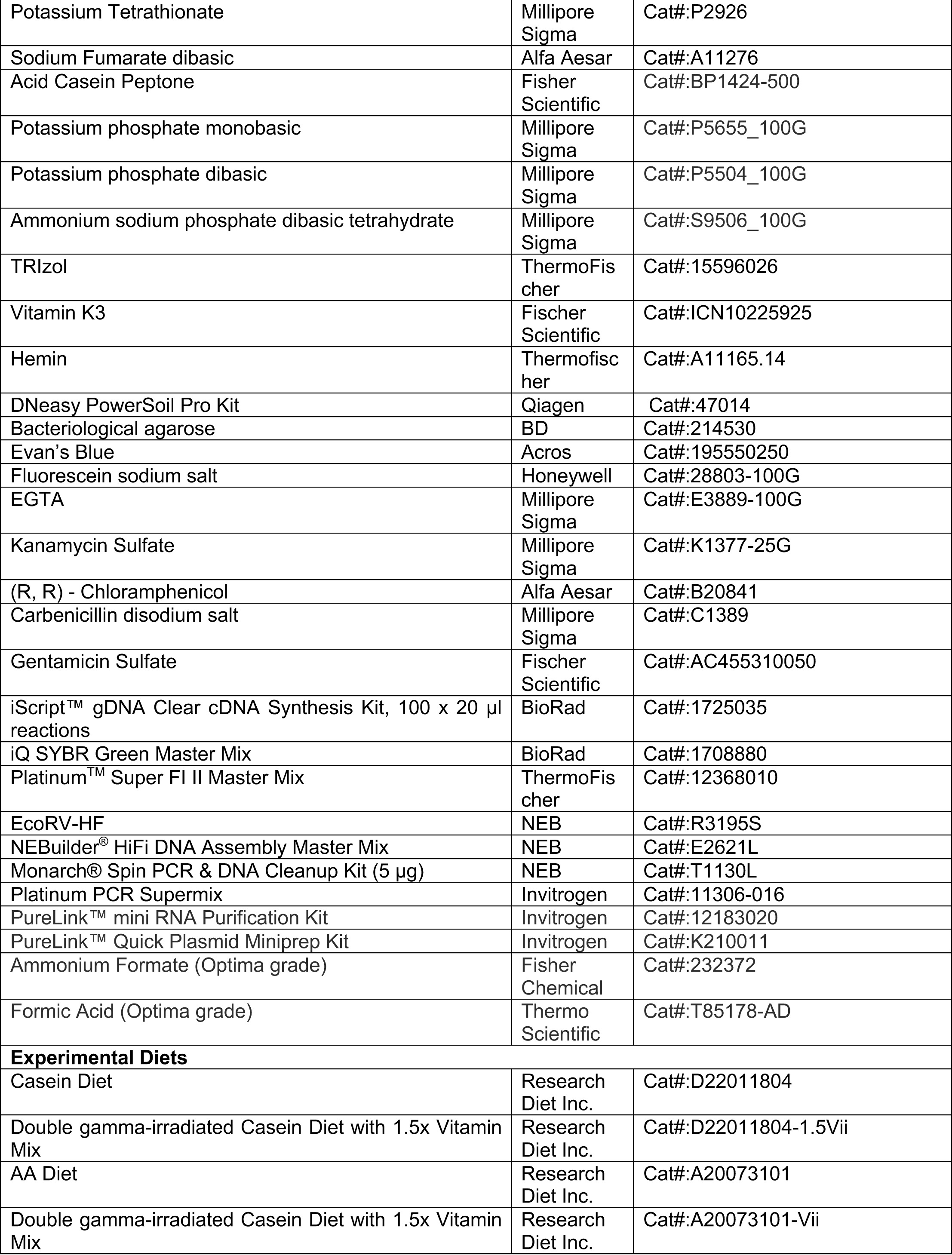
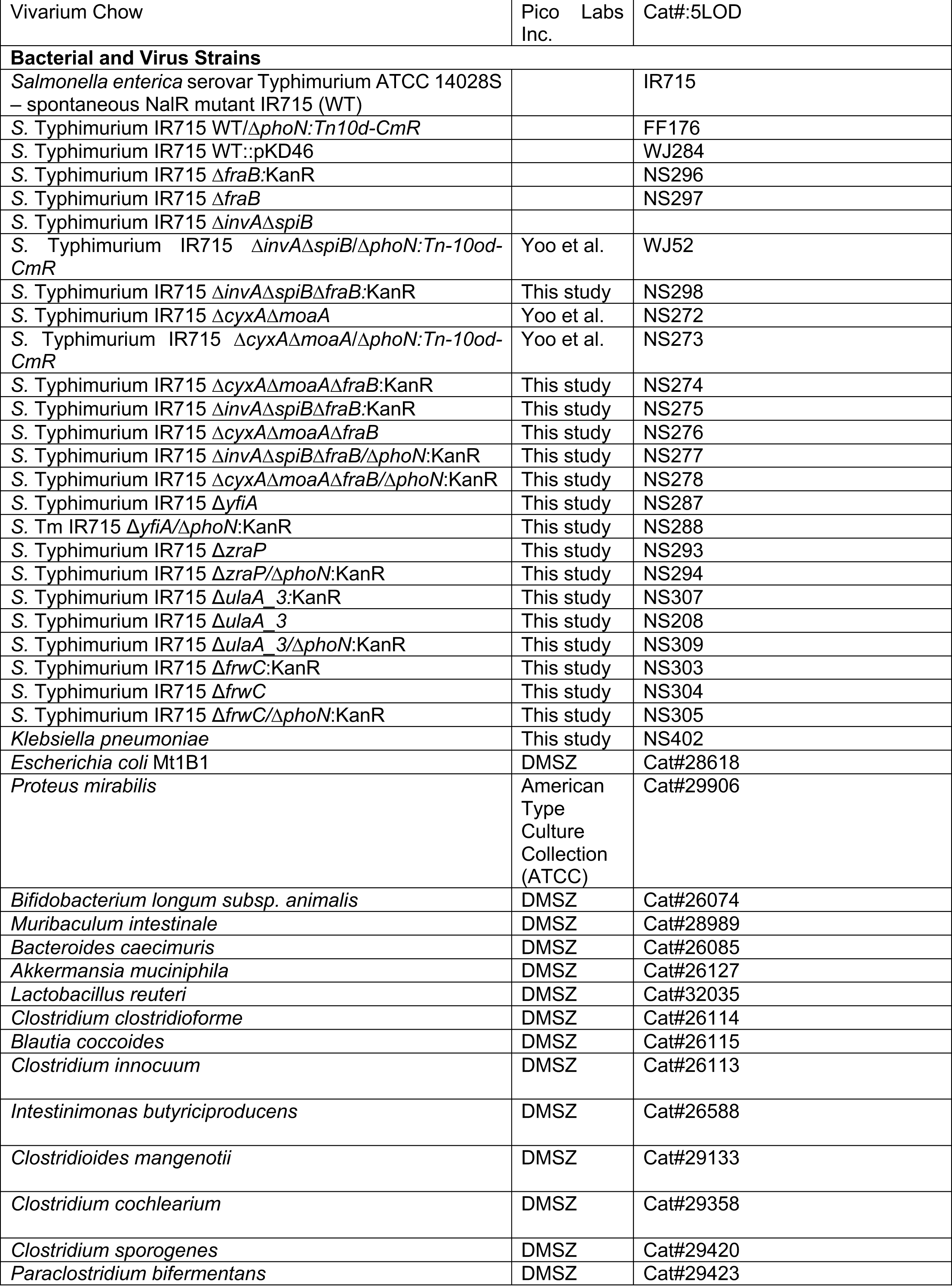
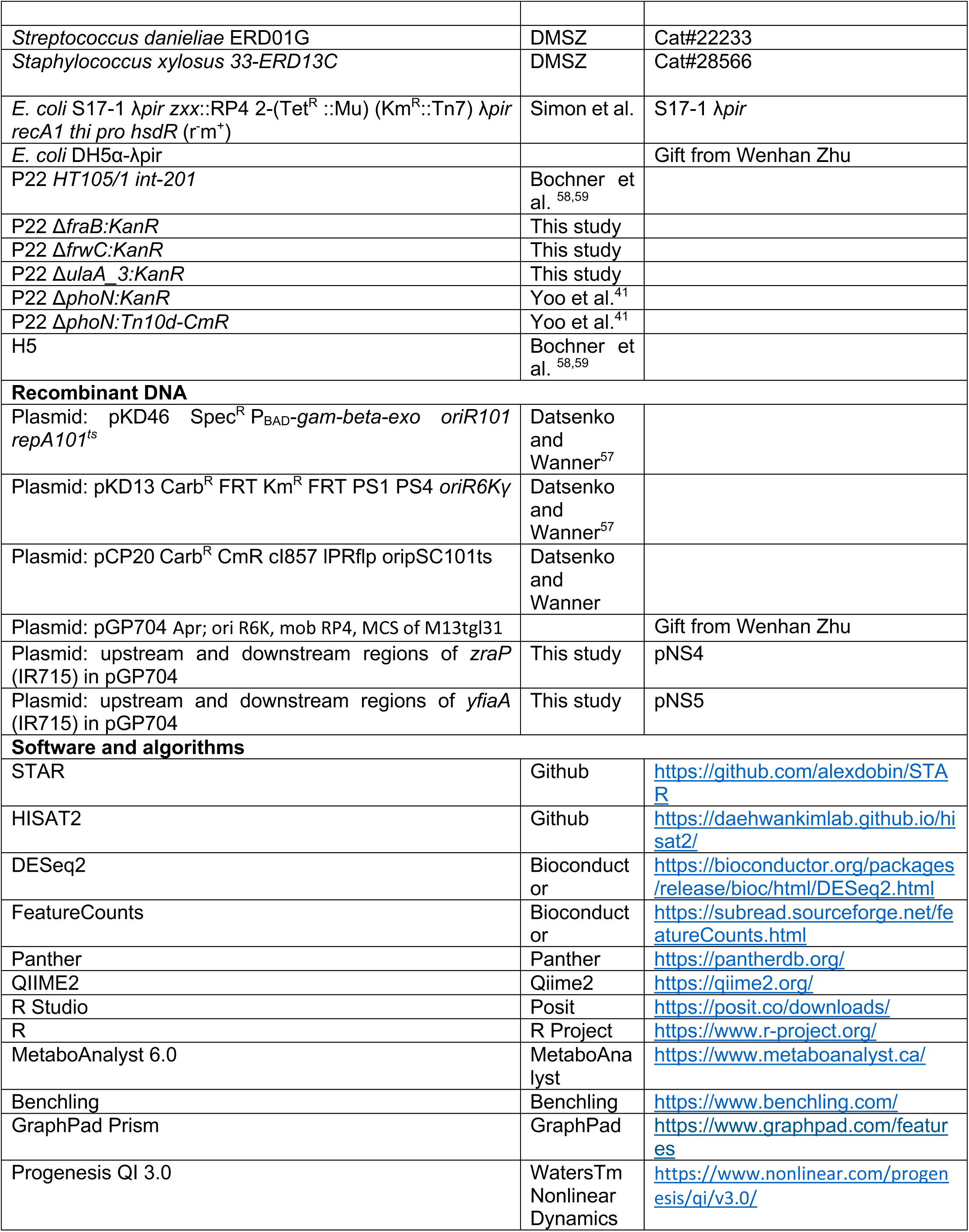

