## Supplementary materials for "Short-term alterations in dietary amino acids override host genetic susceptibility and reveal mechanisms of *Salmonella* Typhimurium small intestine colonization"

**Table S1. Diet formulation of CA (D22011804) and AA diets (A20073101).**

| | **CA Diet** (Cat#:D22011894**)**<br>*Matching Macronutrient Levels to 5L0D* | **AA Diet** (Cat#:A20073101)<br>*Matching Macronutrient Levels to 5L0D* |
|---|---|---|
| **Ingredients** | **grams** | |
| Casein | 280 | 0 |
| L-Cystine | 4.5 | 6 |
| L-Isoleucine | 0 | 10.6 |
| L-Leucine | 0 | 22.2 |
| L-Lysine | 0 | 18.5 |
| L-Methionine | 0 | 7.2 |
| L-Phenylalanine | 0 | 11.8 |
| L-Threonine | 0 | 10 |
| L-Tryptophan | 0 | 3 |
| L-Valine | 0 | 13 |
| L-Histidine | 0 | 6.4 |
| L-Alanine | 0 | 7.1 |
| L-Arginine | 0 | 0 |
| L-Aspartic acid | 0 | 17 |
| L-Glutamic acid | 0 | 53.5 |
| Glycine | 0 | 4.2 |
| L-Proline | 0 | 25 |
| L-Serine | 0 | 14 |
| L-Tyrosine | 0 | 13 |
| **Total L-Amino aicds** | **4.5** | **242.5** |
| Corn Starch | 312 | 312 |
| Maltodextrin 10 | 110 | 110 |
| Dextrose | 150 | 150 |
| Cellulose, BW200 | 100 | 100 |
| Inuline | 46 | 46 |
| Soybean Oil | 61 | 61 |
| Mineral Mix S10026 | 10 | 10 |
| DiCalcium Phosphate | 13 | 13 |
| Calcium Carbonate | 5.5 | 5.5 |
| Potassium Citrate , 1 H2O | 16.5 | 16.5 |
| Sodium Bicarbonate | 7.5 | 7.5 |
| Vitamin Mix V1001 | 10 | 10 |
| Choline Bitartrate | 2 | 2 |
| FD&C Yellow Dye #5 | 0.05 | 0 |
| FD&C Red Dye #40 | 0 | 0.05 |
| **Total** | **1128.05** | **1086.05** |
| | **grams** | |
| **Protein** | 251 | 243 |
| **Carbohydrate** | 599 | 599 |
| **Fat** | 61 | 61 |

| | | |
|---|---|---|
| **Fiber** | 146 | 146 |
| | **grams (%)** | |
| **Protein** | 22.2 | 22.3 |
| **Carbohydrate** | 53.1 | 55.2 |
| **Fat** | 5.4 | 5.6 |
| **Fiber** | 12.9 | 13.4 |
| | **kCal** | |
| **Protein** | 1004 | 970 |
| **Carbohydrate** | 2397 | 2397 |
| **Fat** | 549 | 549 |
| **Fiber** | 3950 | 3916 |
| | **kCal (%)** | |
| **Protein** | 25 | 25 |
| **Carbohydrate** | 61 | 61 |
| **Fat** | 14 | 14 |
| **Fiber** | 100 | 100 |
| **kCal/gram** | **3.5** | **3.6** |

**Table S2. Criteria for blinded scoring of histopathological changes.**

| Score | Submucosal edema | Epithelial damage | Exudate | Submucosal neutrophil infiltration (cells/high power field) | Submucosal mononuclear cell infiltration (cells/high power field) |
|---|---|---|---|---|---|
| 0 | No changes | No changes | No changes | No changes (0-5) | No changes (0-5) |
| 1 | Detectable (<10%) | Desquamation | Slight focal accumulation | (6-20) | (5-10) |
| 2 | Mild (10-20%) | Mild erosion | Mild focal accumulation | (21-60) | (10-20) |
| 3 | Moderate (20-40%) | Marked erosion and/or mild ulceration | Moderate multifocal accumulation | (61-100) | (20-40) |
| 4 | Marked (>40%) | Multifocal ulceration | Marked diffuse accumulation | (>100) | (>40) |

**Table S3. Primers used to construct plasmid or bacterial strains.**

| Name | Sequence |
|---|---|
| zraP_IR715_F1_ERV | CTAGAGGTACCGCATGCGATctcaatatttacagcaaacc |
| zraP_IR715_R1_ERV | tcgcgctaattgccctctctggttatcatcgcggcggcgg |
| zraP_IR715_F2_ERV | ccgccgccgcgatgataaccagagagggcaattagcgcga |
| zraP_IR715_R2_ERV | AATTCCCGGGAGAGCTCGATcatacctactcgcgtgccgg |
| yfiA_IR715_F1_ERV | CTAGAGGTACCGCATGCGATtactataccgcgcgcggcgc |
| yfiA_IR715_R1_ERV | ctggcatctttcaccgatgctgccggagtaatttccattt |
| yfiA_IR715_F2_ERV | aaatggaaattactccggcagcatcggtgaaagatgccag |
| yfiA_IR715_R2_ERV | AATTCCCGGGAGAGCTCGATgagatggtgggctttaccga |
| K1 | CAGTCATAGCCGAATAGCCT |
| FrwC_del_Fwd | TGC CTG GGT GGG TAA CAG CTT CGG CGC GGG TTT TTT CGG G TG TAG GCT GGA GCT GCT TCG |
| FrwC_del_Rev | AAT TAA ACC GCC CCA ACC GGC ATA ACA CTG CGC GCC AAA C ATT CCG GGG ATC CGT CGA CC |
| FrwC_con_Fwd | GTG CGC AGG TTG TGC CAC TC |
| FrwC_con_Rev | GTC GGC GCG TTT TAC CGC GT |
| FraB_del_Fwd | CCT GGA TAA TGG ATT CAC CGT TGG TGG CGC ATT GCG ACT A TG TAG GCT GGA GCT GCT TCG |
| FraB_del_Rev | GAA AAC TGG AAG AAG AAA GGC GTA TTC GCA TCG GTA ATT T ATT CCG GGG ATC CGT CGA CC |
| FraB_con_Fwd | CCG TTT TTG AAT CGG TAG AC |
| FraB_con_Rev | TTC CGA TTC CAG GTG GCC GT |
| ulaA_3_del_Fwd | CAC CAT TAT CTG CAC GGC GAT ACT GGT TTC CCT GTA TG TAG GCT GGA GCT GCT TCG |
| ulaA_3_del_Rev | ACG CAA CCA AAG ATT TCA ATC ATC CCC ATC ACC AGA CAA AATT CCG GGG ATC CGT CGA CC |

| | |
|---|---|
| ulaA_3_con_Fwd | CCA GAT ATC GTG ATG GAA CG |
| ulaA_3_con_Fwd | GAG TGG TTT CAT AGG CGC TA |

**Table S4. qPCR primers used in this study.**

| Name | Sequence |
|---|---|
| Nramp1_qFd | GCAGGCCCAGTTATGGCTC |
| Nramp1_qRv | CAGGCTGAATGTACCCTGGTC |

**Table S5. Plasmids used in this study.**

| | | |
|---|---|---|
| pKD46 $Spec^R$ $P_{BAD}$-*gam-beta-exo oriR101 repA101*$^{ts}$ | Datsenko and Wanner[57] | |
| pKD13 $Carb^R$ FRT $Km^R$ FRT PS1 PS4 *oriR6Kγ* | Datsenko and Wanner[57] | |
| pCP20 $Carb^R$ CmR cI857 lPRflp oripSC101ts | Datsenko and Wanner | |
| pGP704 Apr; ori R6K, mob RP4, MCS of M13tgl31 | | Gift from Wenhan Zhu |
| upstream and downstream regions of *zraP* (IR715) in pGP704 | This study | pNS4 |
| upstream and downstream regions of *yfiaA* (IR715) in pGP704 | This study | pNS5 |

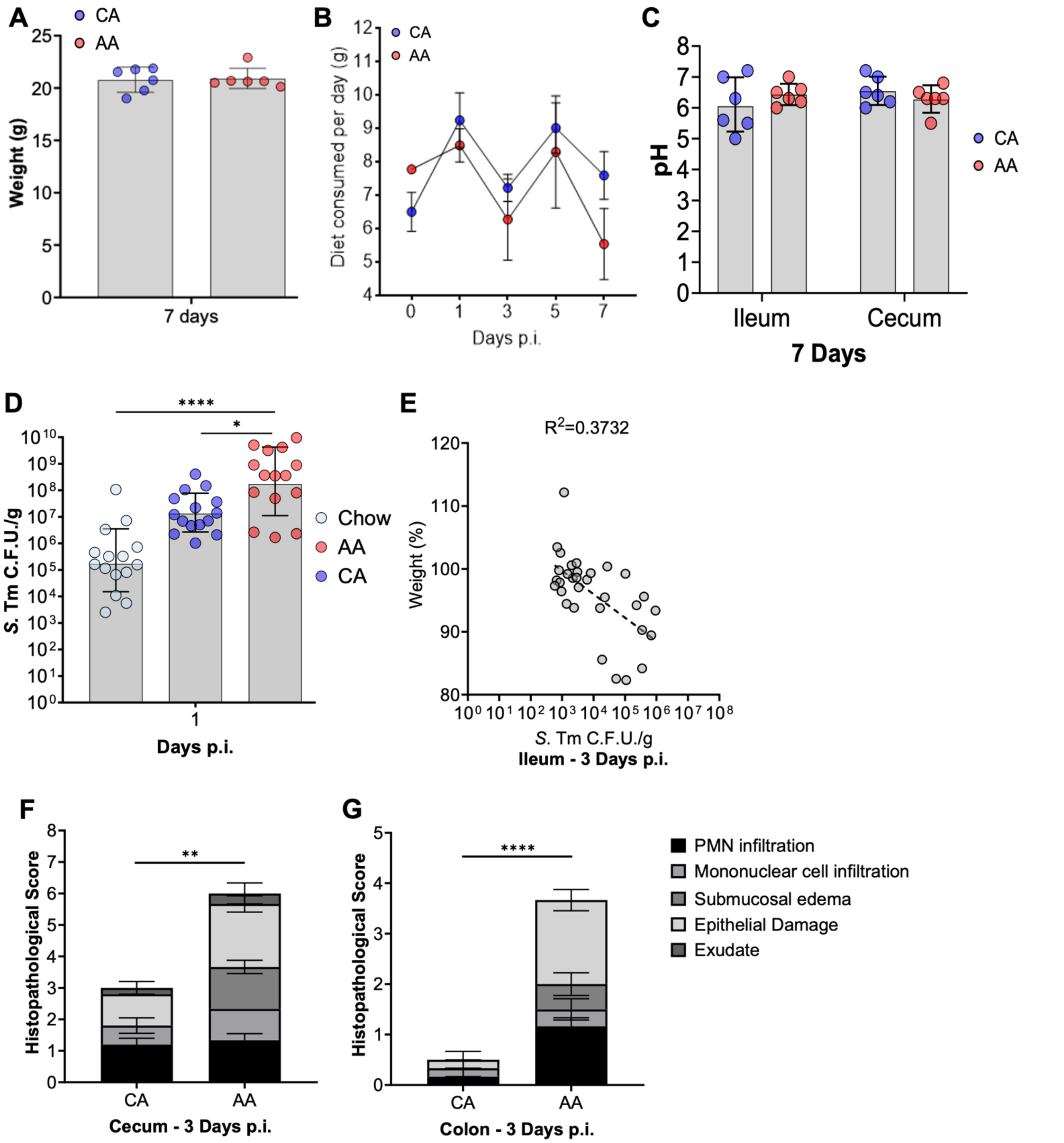


**Figure S1: AA diet alone does not alter mouse weight or luminal pH of the small and large intestines.** Related to Figure 1. (A-C) Naïve CBA/J mice were switched to the CA and AA diet and maintained for 7 days without infection. Every other day, food was weighed to measure the consumption rate over time. At sacrifice, mice were weighed, and intestinal content was harvested to measure pH. (D) *S*. Tm burden in feces one d.p.i. of Chow, CA, or AA diet-fed mice. Geomean and geometric SD, ANOVA, P=*,<0.05; ****,<0.0001.(E) Correlation between weight loss and bacterial burden in the terminal ileum of mice. (F-G) Histopathological scoring of cecal and proximal colon tissue three d.p.i. between CA and AA diet-fed mice. Mann Whitney, P=**,<0.01; ***,<0.001.

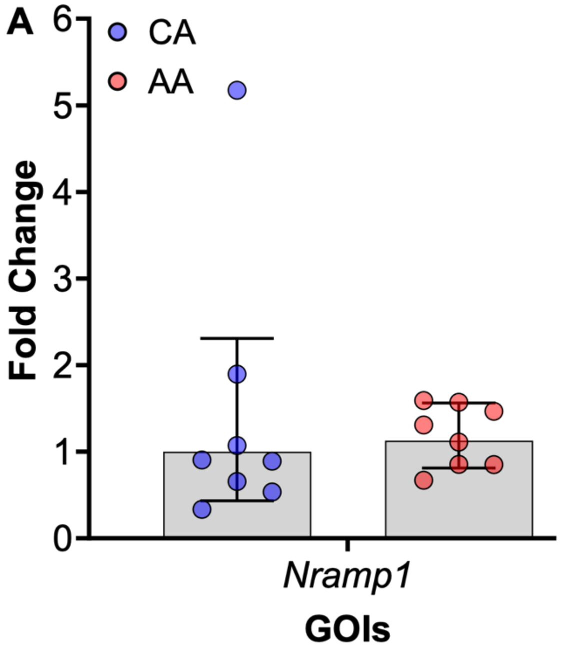


**Figure S2: AA diet-fed demonstrate no differences in *Nramp1* expression 3 d.p.i. in the ileum compared to the CA diet-fed mice.** Related to the Figure 2. (A) CBA/J mice were either maintained on Chow or switched to a CA or AA diet two days before infection with *S*. Tm. *Nramp1* expression in the ileum of mice measured by RT-qPCR three d.p.i.. n=8.

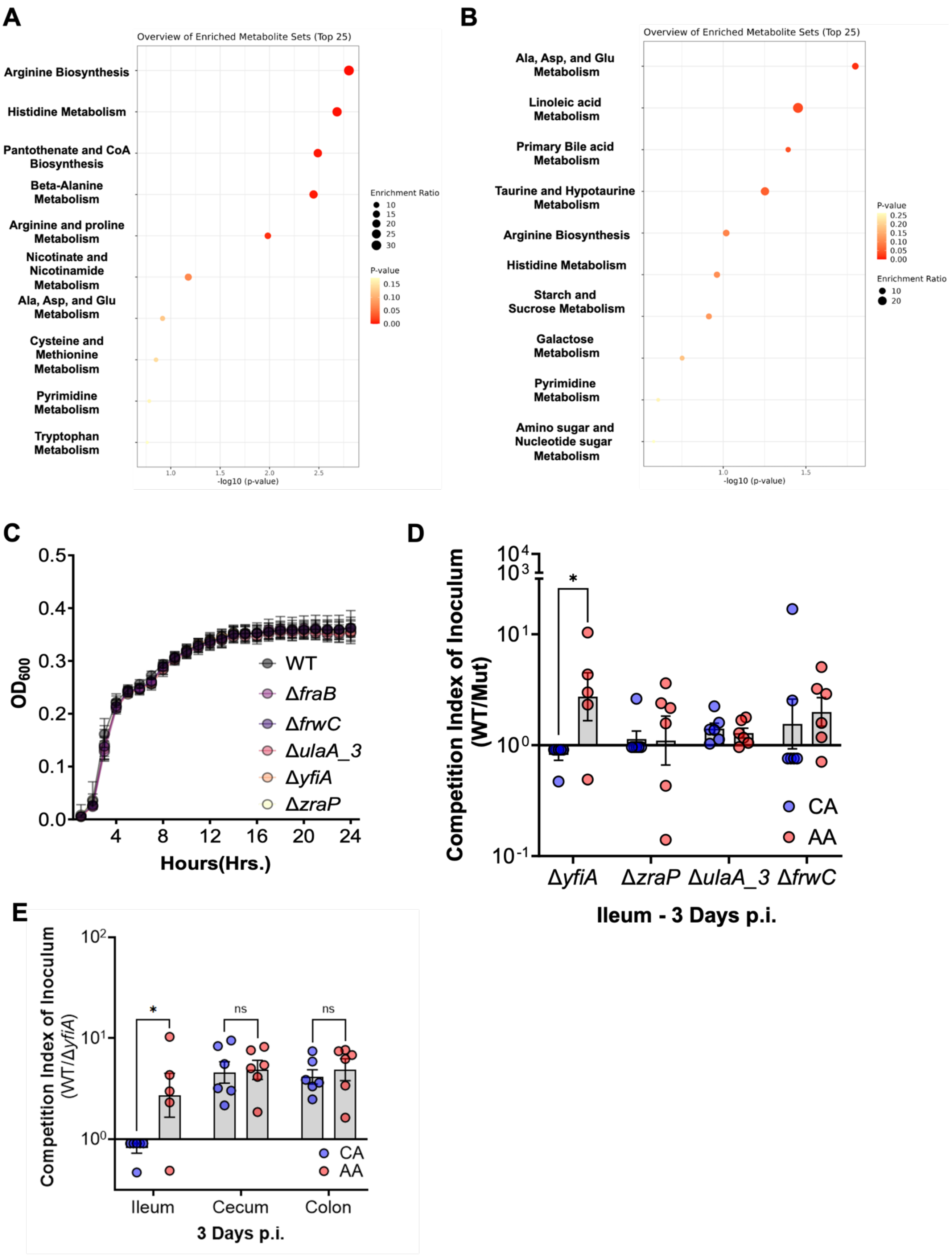


**Figure S3: AA diet alters the metabolic profile of the ileum overall before infection and alters *S.* Tm metabolism *in vivo*.** Related to Figure 6. Metabolites significantly altered (P<0.05 PERMANOVA; fold change greater than 2) were used for enrichment analysis via Metaboanalyst 5.0. (A) Enriched in AA-diet-fed mice; (B)

Depleted in AA-diet-fed mice. (C-D) To interrogate the relationship between our RNAseq and Metabolomics data were constructed five in-frame deletions associated with the top five DEGs or pathways within *S*. Tm. (C) Strains cultured in biological triplicate anaerobically in LB *in vitro*, measuring optical density at 600nm every hour for up 24 hours. (D) CBA/J mice pre-fed either a CA or AA diet infected with a 1:1 competitive ratio of wild type and an isogenic strain deficient in *yfiA, zraP, ulaA_3*, or *frwC;* and allowed carriage for three d.p.i.. competition index of inoculum in the ileal lumen. (E) Competitive index of mice infected with a 1:1 competitive ratio of wild type and isogenic mutants deficient in *yfiA* in the ileum, cecum, and colon content of CA- and AA diet-fed mice. N=6 per diet group. Student's t-test or Multiple t-test, P=*,<0.05.
